# Dual-specificity phosphatases 13 and 27 as key switches in muscle stem cell transition from proliferation to differentiation

**DOI:** 10.1101/2023.12.25.570044

**Authors:** Takuto Hayashi, Shunya Sadaki, Ryosuke Tsuji, Risa Okada, Sayaka Fuseya, Maho Kanai, Ayano Nakamura, Yui Okamura, Masafumi Muratani, Gu Wenchao, Takehito Sugasawa, Seiya Mizuno, Eiji Warabi, Takashi Kudo, Satoru Takahashi, Ryo Fujita

## Abstract

Muscle regeneration depends on muscle stem cell (MuSC) activity. Myogenic regulatory factors, including myoblast determination protein 1 (MyoD), regulate the fate transition of MuSCs. However, the direct target of MYOD in the process is not completely clear. Using previously established MyoD knock-in (MyoD-KI) mice, we revealed that MyoD targets dual-specificity phosphatase (Dusp) 13 and Dusp27. In *Dusp13*:*Dusp27* double knock-out (DKO) mice, the ability for muscle regeneration after injury was reduced. Moreover, single-cell RNA sequencing of MyoD-high expressing MuSCs from MyoD-KI mice revealed that *Dusp13* and *Dusp27* are expressed only in specific populations within MyoD-high MuSCs, which also express *Myogenin*. Overexpressing *Dusp13* in MuSCs causes premature muscle differentiation. Thus, we propose a model where DUSP13 and DUSP27 contribute to the fate transition of MuSCs from proliferation to differentiation during myogenesis.

**Significance Statement:** MYOD protein is not expressed in quiescent muscle stem cells but accumulates rapidly following muscle injury, leading to the proliferation of myogenic progenitors for differentiation. However, the direct targets of MYOD, aside from myogenin, which play roles in myogenic differentiation remain incompletely understood. Using previously established MyoD knock-in mice and single-cell RNA sequencing, we discovered that Dusp13 and Dusp27 are potential target genes of MYOD that promote myogenesis during muscle regeneration in adult mice.

## Introduction

Skeletal muscle stem cells (MuSCs) [1] play a vital role in postnatal muscle development and skeletal muscle regeneration [2]. MuSCs are located near the myofibers beneath the basal lamina and exist in a quiescent state in healthy muscle under resting conditions. However, in response to injury, they become activated, differentiate, and generate new myofibers [3,4]. Some of the progeny of the activated MuSCs undergo self-renewal to perpetually sustain the MuSC pool throughout a lifespan [5–7].

The dynamics of MuSCs are regulated intricately by a series of myogenic regulatory factors (MRFs) that are expressed sequentially during myogenesis. Quiescent MuSCs express the paired homeodomain family transcription factor PAX7 [8,9]. Loss of Pax7 results in normal embryonic and fetal myogenesis but reduced postnatal muscle growth [9]. Interestingly, MuSCs are present in Pax7 mutants, although the population decreases progressively as mice mature due to cell cycle defects and apoptosis [10,11]. The MRFs, including MyoD, Myf5, myogenin, and MRF4, act as basic helix–loop– helix (bHLH) transcription factors that bind to E-box sites. MYOD and MYF5 are essential transcription factors for muscle lineage specification [12,13], MYOGENIN and MRF4 are indispensable for muscle differentiation [14,15]. Additionally, MRFs are induced in activated MuSCs during muscle regeneration in adult mice. Quiescent MuSCs repress the translation of specific transcripts that keep them primed to rapidly activate the myogenic program in case of injury [16]. MicroRNAs and RNA-binding proteins primarily mediate the repression of *Myf5* and *MyoD* transcripts [17–20]. Upon injury, the block on translation is lifted, leading to the rapid translation of *MyoD* and *Myf5*, initiating the myogenic program for regeneration.

*MyoD* KO mice exhibited delayed myogenic differentiation and accelerated self-renewal ability in MuSCs. Furthermore, *MyoD* KO mice on an mdx background, a model for Duchenne muscular dystrophy, have a low survival rate [21,22] . MYOD plays a critical role in determining the fate of activated MuSCs by balancing self-renewal and differentiation during myogenesis. However, the direct targets of MYOD, apart from *myogenin* that play critical roles in myogenic differentiation, are not completely understood.

In a prior study, we established a MyoD knock-in (KI) reporter mouse line to recapitulate the expression levels of endogenous MyoD using tdTomato fluorescence [23]. The fluorescence allowed us to identify the status of MuSCs during myogenesis. We observed that the MyoD-low population, characterized by low tdTomato fluorescence, consisted mostly of undifferentiated stem-like cells. Conversely, the MyoD-high population, with high tdTomato fluorescence, exhibited features of committed and differentiating MuSC [23]. To investigate the molecular mechanisms governing the stemness and commitment/differentiation of MuSCs, whole-transcriptome data were collected from MyoD-low MuSCs, MyoD-high MuSCs, and quiescent MuSCs isolated from Pax7^YFP/YFP^ (Pax7-YFP) mice [23,24]. The establishment of MyoD-KI mice and the availability of whole-transcriptome data at different activation stages of MuSCs based on MYOD levels provide a valuable platform for understanding the fate determination mediated by MYOD.

The DUSP family comprises members of the protein tyrosine phosphatase superfamily involved in the dephosphorylation of phosphotyrosine and phosphoserine/threonine residues [25–27]. They interact directly with MAPK effectors, including extracellular signal-regulated kinases 1/2 (ERK1/2), leading to their dephosphorylation and subsequent inactivation. Therefore, DUSPs play regulatory roles in cell growth, cell differentiation, and apoptosis [28–30]. The phosphatase domain common to DUSPs is known as protein tyrosine phosphatases (PTP). DUSPs can be classified into two subfamilies based on the presence or absence of the MAP kinase-binding (MKB) motif or the kinase-interacting motif (KIM). The DUSPs with the KIM domain are classified as typical DUSPs, whereas those lacking the KIM domain are classified as atypical DUSPs [31,32].

The functional regulation of substrates by DUSPs does not always require phosphatase activity. DUSPs can control the function of MAPKs by sequestering them in the cytoplasm or nucleus. It has also been observed that some DUSPs are catalytically inactive [33]. For example, DUSP27 lacks phosphatase activity due to the absence of a highly conserved cysteine residue essential for enzyme activity [34]. Nevertheless, the loss of *Dusp27* in zebrafish leads to severe disorganization of the sarcomere structure, underscoring the significance of its pseudophosphatase function in muscle development [34]. Within skeletal muscles, DUSP5 and DUSP6, classified as typical DUSPs, regulate muscle hypertrophy and myofiber types by inhibiting ERK1/2 signaling [35]. However, our understanding of the function of atypical DUSPs in skeletal muscle in mammals is limited.

In the present study, we used our MyoD-KI transcriptome and various datasets to identify *Dusp13* as a direct target for MYOD. *Dusp13* is located on chromosome 10a22.2 and encodes two distinct DUSPs, *Dusp13a* and *Dusp13b*. Although DUSP13a is expressed predominantly in murine skeletal muscles [36], its specific roles in skeletal muscle remain unclear. Single KO of *Dusp13* did not exhibit any regenerative defects; however, there was compensatory upregulation of *Dusp27* in skeletal muscles lacking *Dusp13*. To further investigate, we generated a double KO (DKO) of *Dusp13* and *Dusp27* and observed diminished muscle regeneration activity after acute muscle injury. Furthermore, through single-cell RNA-seq (scRNA-seq) analysis, we found that the expressions of *Dusp13* and *Dusp27* are limited to specific subsets of MyoD-high expressing MuSCs that also show the expression of late myogenic markers. Based on the findings, we propose a model in which DUSP13 and DUSP27 instigate the transition of MuSCs from an activated/proliferative state to a differentiation state during regeneration.

## Materials and Methods

### Mice

Mice were kept in a specific pathogen-free environment at the Laboratory Animal Resource Center at the University of Tsukuba, Ibaraki, Japan. All experiments were performed in compliance with the relevant Japanese and institutional laws and guidelines and were approved by the University of Tsukuba Animal Ethics Committee (authorization number: 23-046). The study used 10–15-week-old C57BL/6 J male and female WT, *Dusp13* KO, *Dusp27* KO, and *Dusp13*/*Dusp27* DKO mice, unless otherwise stated. Quiescent MuSCs were isolated directly from the leg muscles of 15–16-week-old male Pax7-YFP (*Pax7*^YFP/YFP^) [24] [23]. The TA muscle was acutely injured by injecting 50 μL of 10 µM cardiotoxin (Funakoshi Co., Ltd., Tokyo, Japan) under anesthesia using isoflurane inhalation.

### Generation of *Dusp13* and *Dusp27* KO mice

The *Dusp13* and *Dusp27* KO mice were generated using the CRISPR/Cas9 system. Two target sgRNAs were designed as follows; 5 ′ -GTAGGGTTCACACAGAACGG-3 ′ and 5 ′ - ATGTCACAGTAAGCGTACGG-3′ for targeting all exons of the *Dusp13* gene on mouse chromosome 14, and 5 ′ -GCATCCTTCCACATCAACCC-3 ′ and 5 ′ - GGTAAGGGTGCTCTCGACAA-3′ for targeting regions located in introns 2 and 3, respectively, of the *Dusp27* gene on mouse chromosome 1. Each target sgRNA was cloned into the guide RNA-Cas co-expression plasmid px330-U6-Chimeric_BB-CBh-hSpCas9 vector (Addgene, Watertown, MA, USA). A mixture of the four cloned plasmids was microinjected into the pronuclei of C57BL/6 J fertilized eggs, and living embryos were transferred to pseudo-pregnant ICR female mice. Genomic DNA screening was performed using PCR with the primers and protocols listed in Table S1.

### Muscle stem cell isolation and culture

MuSCs were isolated from the hindlimb muscles of 11–13-week-old mice following MACS using a Satellite Cell Isolation Kit (Miltenyi Biotec, Bergisch Gladbach, Germany) and anti-integrin α7 microbeads (Miltenyi Biotec), as previously described [23]. The adult hindlimb muscles were minced in ice-cold Ham’s F12 medium. The minced muscles were then incubated in 0.1% collagenase D (Roche, Basel, Switzerland) and 0.1% trypsin in Ham’s F12 (Gibco, Waltham, MA, USA) with 1% penicillin-streptomycin for 45 min at 37 °C on a shaker for three rounds of digestion. After each round, the supernatant was collected in a 50 mL tube containing 8 mL of fetal bovine serum (FBS) (Gibco) on ice, and fresh 0.1% trypsin/0.1% collagenase digestion buffer was added to the tube for the next round of digestion. The collected supernatant was filtered through a sequence of 100 and 40 μm nylon mesh strainers and then centrifuged at 400 × *g* for 10 min at 4 °C. The pellets were resuspended in PBS containing 2% FBS and processed for MACS or flow cytometry. The MuSCs were cultured in a growth medium containing 39% DMEM (Gibco), 39% Ham’s F12, 20% FBS (Gibco), and 1% UltroserG (Pall Life Sciences, Port Washington, NY, USA). Differentiation media consisted of 2% horse serum in Dulbecco’s modified Eagle’s medium (DMEM). Gelatin (0.2%)-coated dishes were used for MuSC culture unless otherwise indicated.

### Flow cytometry analysis and cell sorting

MACS-isolated MuSCs were cultured for 5 days. They were detached using 0.25% trypsin/EDTA, then centrifuged and resuspended in 2% FBS/PBS. Propidium iodide (PI) was added to the cells at a 1:500 (vol/vol) ratio and incubated on ice for 10 min. The cells were either analyzed using a flow cytometer (CytoFLEX S; Beckman Coulter, Brea, CA, USA) or sorted using a MoFlo XDP flow cytometer (Beckman Coulter). Debris and dead cells were excluded using forward scatter, side scatter, and PI gating. Gates were defined based on WT control fluorescence. The data were analyzed using FlowJo^TM^ software ver. 10.7.1 (BD Biosciences, Franklin Lakes, NJ, USA).

### Immunofluorescence and histological analysis

Cultured cells were washed twice with PBS after removing the media and fixed with 4% paraformaldehyde (PFA) for 15 min. The fixed cells were blocked with 2% BSA/PBS for 1 h at room temperature and then washed three times with PBS. Isolated single EDL myofibers [6] were fixed using 2% PFA for 15 min, permeabilized with 0.1% Triton in PBS, and blocked with 5% goat serum in 0.1% Triton/PBS for 30 min on a shaker. The cultured cells and single EDL myofibers were incubated with primary antibodies at 4 °C overnight. The myotube diameter and nucleus number were measured using the ImageJ software (NIH, Bethesda, MI, USA). The myotube diameter was defined as the average diameter at the three ends (both ends and the center) for each MYHC-positive and multinucleated myotube. TA muscles were neatly stacked on top of each other. The Achilles tendon side was vertically mounted on tragacanth gum on a cork disc and quickly frozen in isopentane cooled in liquid nitrogen. Thin frozen muscle sections (8 µm) were subjected to immunohistochemical analyses. The sections were fixed with 4% PFA for 10 min at room temperature. For PAX7 staining, sections underwent antigen retrieval with DAKO buffer (Agilent, Santa Clara, CA, USA) for 10 min at 110 °C, followed by blocking with M.O.M. blocking reagent (Vector Laboratories, Newark, CA, USA). The sections were then incubated with primary antibodies overnight at 4 °C. Tissue sections were mounted with VECTASHIELD Vibrance Antifade Mounting Medium with DAPI (Vector Laboratories). Primary antibodies were against PAX7 (Developmental Studies Hybridoma Bank (DSHB)), MYOD (sc-377460; SantaCruz), MYOD (ab133627, abcam), RFP (600-401-379, Rockland), LAMININ (sc-59854; SantaCruz), MYOGENIN (124800; Abcam, Cambridge, UK), MYOGENIN (MA5-11486; Invitrogen, Waltham, MA, USA), (MyHC (DSHB; MF20), and KI67 (MA5-14520; Invitrogen, Waltham, MA, USA). For immunofluorescence, secondary antibodies conjugated with Alexa Fluor-488, Alexa Fluor-546, Alexa Fluor-555, Alexa Fluor-633, and Alexa Fluor-647 were used. The secondary antibodies included anti-mouse IgG1, anti-mouse IgG2b, anti-rat IgG, and anti-rabbit antibodies (Thermo Fisher Scientific, Waltham, MA, USA). For HE staining, the SOL, PLA, and TA muscle sections were stained with Mayer’s hematoxylin (Wako, Richmond, VA, USA) for 10 min. After washing in warm running tap water (50 ± 5 °C) for 3 min, eosin (Wako) was applied for 1 min. The sections were then dehydrated using two changes (5 s each) of 95% ethanol and two changes (5 s each) of 100% ethanol (Wako), followed by three changes of xylene (Wako) for 5 min each. Finally, the sections were mounted with DPX mounting medium (Sigma-Aldrich, St. Louis, MA, USA) and observed with a BIOREVO BZ-X800 microscope (Keyence, Osaka, Japan). The CSA of myofibers was assessed using the same microscope system and a hybrid cell count application (Keyence).

### RNA-scope

Male Pax7-YFP mice, aged 12–15 weeks, were used to directly isolate quiescent MuSCs from the leg muscles. A total of 20,000 sorted cells were seeded in a well of a 24-well culture plate, pre-coated with iMatrix-511 (Nippi) for 1 h, and immediately fixed with 4% paraformaldehyde (PFA) for 15 min (freshly isolated MuSCs). The sorted cells were cultured in growth medium for 6 days (GM6) or in differentiation medium for 6 days (DM6) and then fixed with 4% PFA for 15 min. For the isolation of MyoD-low and MyoD-high MuSCs, MACS-isolated MuSCs from MyoD-KI mice were cultured for 5 days, detached using 0.25% trypsin/EDTA, followed by centrifugation and resuspension in 2% FBS/PBS, as previously described [23]. The cells were sorted using a MoFlo XDP flow cytometer (Beckman Coulter), and dead cells were excluded by propidium iodide (PI) gating. The sorted MyoD-low and MyoD-high MuSCs were seeded in a well of a 24-well culture plate pre-coated with iMatrix-511 (Nippi) for 30 min, then immediately fixed with 4% PFA for 15 min.

For the detection of Dusp13 and Dusp27 transcripts, the fixed cells were processed using the RNAscope™ Multiplex Fluorescent V2 Assay (ADC, 323270). Dusp13 and Dusp27 transcripts were hybridized with Mn-Dusp13a-O1-C1 (ADC, 1332391-C1) and Mn-Styxl2 (Dusp27)-C1 (ADC, 1332401-C1) probes, respectively. Briefly, the fixed cells were washed twice with PBS. After a 10-minute incubation in hydrogen peroxide solution, the cells were incubated with protease III diluted 1:15 in PBS for 10 min. Following two PBS washes, the cells were hybridized with each C1 probe for 2 h at 40°C. After the amplification steps in the kit, Dusp13 and Dusp27 transcripts were visualized using the TSA Vivid Fluorophore Kit 520 (R&D Systems, 7523). Finally, the cells were washed twice with PBS, and subsequent immunofluorescence analysis was performed using antibodies against PAX7, MYOD, MF20, and RFP, as described above. Nuclei were counterstained with DAPI.

### DNA construct

The *MyoD*, *Dusp13*, and *Dusp27* genes were amplified using PrimeStar GXL DNA Polymerase (Takara Bio, Shiga, Japan) with soleus cDNA as the template. After confirming the size of the PCR products using 1% agarose gel electrophoresis, the target PCR product was purified from the gel. The purified DNA fragment of *Dusp13* was digested with EcoRI. The digested fragments were then ligated to pENTR3c-IRES-EGFP, which was also digested with the same restriction enzymes, using Ligation High (TOYOBO, Osaka, Japan). DUSP13 deficient in phosphatase activity (Dusp13_Mut) was generated using the PrimeSTAR® Mutagenesis Basal Kit (Takara Bio). The Dusp family contains the amino acid cysteine, which is essential for enzyme activity [37]. The present mutant was generated using Dusp13_D97A and Dusp13_C128S primers (Table S1) to replace aspartic acid at position 97 and cysteine at position 128 with alanine and serine, respectively. The reaction was then transformed into One shot TOP10 Competent cells (Thermo Fisher Scientific) and seeded on kanamycin-LB agar plates. A single colony was grown in kanamycin-LB medium, and after confirming each insert by colony PCR and DNA sequencing, plasmid purification was performed using the FastGene Plasmid Mini Kit. Clones without any mutations in the product were selected. The LR reaction of the Gateway system was performed to recombine pENTR3c-Dusp13-IRES-EGFP into pAd/CMV/V5-DEST (Thermo Fisher Scientific). In total, 100 ng of the purified plasmid and 300 ng of pAd/CMV/V5-DEST were mixed at room temperature and reacted according to the manual for Gateway LR Clonase II Enzyme Mix (Thermo Fisher Scientific). The 5 μL reaction was subsequently transformed into 50 μL of competent cells and seeded on ampicillin-LB agar plates. Single colonies were cultivated in LB medium, and confirmation of the insert was achieved through colony PCR. The confirmed plasmid was prepared in large quantities using the PureLink HiPure Plasmid Midiprep Kit (Thermo Fisher Scientific). The plasmid concentration was measured using a NanoDrop 2000 Spectrophotometer (Thermo Fisher Scientific), and 20 μg of the plasmid was digested with the restriction enzyme PacI. Verification of complete digestion by PacI was performed through 1% agarose gel electrophoresis, and the linearized plasmid was purified via phenol:chloroform extraction and ethanol precipitation. To generate the luciferase vector, the purified DNA fragment of the *MyoD* gene was cloned into a pAAV-CMV vector using the In-Fusion Cloning HD Kit (Takara Bio). The mouse *Pax7 and Myogenin* sequence was amplified by PrimeSTAR^®^ Max DNA polymerase (TAKARA) using the primers listed in the Table S1. The purified *Pax7* fragment was cloned into *Bam*HI-*Sal*I site of pLenti-PGK-GFP [38] using In-Fusion cloning HD kit (TAKARA). The purified *Myogenin* fragment was cloned into *Xho I* site of pcDNA-EGFP (gift from Doung Golenbock, Addgene plasmid #13031) using In-Fusion cloning HD kit (TAKARA). The DNA fragment corresponding to the 300 bp 5′ upstream region (−250 to +50) of *Dusp13* or the 1,200 bp upstream region (−1,000 to +200) of *Dusp27* from EDL muscle DNA of C57BL/6 mice was inserted into a pGL4.10 promoter-less luciferase vector (Promega, Madison, WI, USA) digested with XhoI. The resulting constructs were named pGL4-Dusp13 or pGL4-Dusp27. The primer sequences for the construction of vectors are provided in Table S1.

### Adenovirus production and infection

293A cells were seeded on BD BioCoat Collagen I 60 mm Culture Dishes (IWAKI) using the medium (DMEM and 10% FBS). Purified 5 μg of pAd/CMV/V5-DEST plasmid was transfected using Fugene HD (Promega) when the 293A cells were 80% confluent. When the cells began to float, the bottom surface of the dish was scraped off using a scraper and collected in a 15 mL High-Clarity Polypropylene Conical Tube 17 × 120 mm style (FALCON) (primary virus). The primary virus underwent repeated freeze-thaw cycles in a water bath at 37 °C and in liquid nitrogen. This freeze-thaw process was repeated three times. Finally, it was frozen again and stored at −80 °C. Next, 2 mL of the primary virus was thawed at 37 °C and centrifuged for 15 min at 3,000 rpm (KUBOTA 2800). The supernatant was added to 293A cells (80% confluent) cultured in a 100 mm Collagen-Coated Dish Collagen I (IWAKI) and infected for 1 h at 37 °C in a CO_2_ incubator. Then, 8 mL of medium was added. When the cells floated, they were scraped off using a scraper and collected in 15-mL tubes. The freeze-thaw process was repeated three times. Finally, it was frozen again and stored at −80 °C. This virus was created as the 2nd virus. This procedure was repeated five times, and the 5th virus was stored at −80 °C. Next, to purify the virus, the fifth virus was freeze-thawed three times and ultracentrifuged at 23 k rpm at 4 °C for 1.5 h in the Super-centrifuge Optima XE-90 Ultracentrifuge (Beckman Coulter) using Ultra-Clear Centrifuge Tubes (Beckman Coulter). After ultracentrifugation, recombinant adenovirus was extracted from the second viral layer from the bottom. The dialysis buffer was prepared in three heat-resistant plastic bottles by adding 1976 mL of MilliQ, 20 mL of 1 M Tris HCl (pH 8.0), and 4 mL of 1 M MgCl_2_, and the bottles were then autoclaved. For dialysis, the recombinant adenovirus extract was injected into Slide-A-Lyzer Dialysis Cassettes (Thermo Fisher Scientific) and dialyzed in a dialysis buffer at 4 °C with stirring. The dialysis buffer was changed three times every 12 h. The purified recombinant adenovirus was stored at −80 °C. Infectious titers were measured using the Adeno-X Rapid Titer Kit (Takara Bio). After 48 h from the start of differentiation, the medium was replaced with a fresh DM and incubated with adenovirus at a multiplicity of infection (MOI) of 250 in a differentiation medium. Two days after the virus transduction, the cells were fixed with 4% PFA/PBS.

### Luciferase assay

To determine if MyoD, Pax7 and Myogenin could activate the *Dusp13* or *Dusp27* promoter, we co-transfected 100 ng of pGL4-Dusp13 or pGL4-Dusp27 with 50 ng of pRL-TK vectors (Promega) and varying amounts (0, 10, 20, 100, 200 ng) of MyoD, Pax7, and Myogenin expression vectors into 293T cells (Takara Bio) using the FuGENE HD transfection reagent (Promega). Twenty-four hours after transfection, the cells were harvested and assayed using the Dual-Luciferase Reporter Assay System (Promega). Luciferase activity was measured using the GloMax® 20/20 Luminometer (Promega) and normalized based on Renilla luciferase activity from the pRL-TK vector.

### Motif enrichment analysis

The presence of Ebox sites in the promoter regions of genes was predicted using the Find Individual Motif Occurrences (FIMO) algorithm from the MEME Suite [39], utilizing the position weight matrix available from JASPAR (https://jaspar.genereg.net/). This analysis was conducted from 1 kbp upstream to 500 bp downstream of the transcriptional start site of *Dusp13*/*Dusp27*.

### RNA isolation and RT-qPCR

Total RNA was extracted from various tissues and MuSCs using ISOGEN (NIPPON GENE, Chiyoda, Tokyo, Japan). We synthesized first-strand cDNA using the QuantiTect Reverse Transcription Kit (Qiagen, Hilden, Germany). Polymerase chain reaction (PCR) was then performed using a THUNDERBIRD SYBR qPCR system (TOYOBO) and a TP850 Thermal Cycler Dice Real Time System (Takara Bio, Shiga, Japan). The primer sequences are listed in Table S1. The relative amount of each transcript was normalized to the number of beta-actin transcripts.

### Single-cell RNA-seq analysis

MACS-isolated MuSCs from MyoD-KI mice [23] were cultured for five days and then detached using 0.25% trypsin/EDTA. The cells were centrifuged and resuspended in 2% FBS/PBS. The cells were incubated with PI at a 1:500 (v/v) ratio. Debris and dead cells were excluded using forward scatter, side scatter, and PI gating. Live MyoD-tdTomato-high MuSCs were sorted using a MoFlo XDP flow cytometer (Beckman Coulter) based on our previous study’s gate of MyoD-tdTomato-high population. Single RNA-seq libraries from MyoD-tdTomato-high MuSCs were generated using a chromium controller (10× Genomics, Pleasanton, CA, USA) following the manufacturer’s instructions. Approximately 10,000 cells were processed using the Chromium Next GEN Single Cell 3’Reagent Kits v3.1 (10× Genomics). The sequencing was performed on a NovaSeq 6000 (Illumina, San Diego, CA, USA) operated by the Rhelixa Corporation (Tokyo, Japan). The fastq files were processed using default parameters for adaptor sequence filtering, and low-quality reads were removed to obtain clean data. Feature-barcode matrices were then generated by aligning the reads to the mouse genome (mm10 Ensemble: version 92) with CellRanger v3.1.0. The scRNA-seq data was processed using the standard Seurat pipeline. For quality control, the range of gene reads per cell was set between 300 and 6000. Additionally, the filtering threshold for mitochondrial gene proportion was established at 15%. The “FindVariableFeature” function was used to determine the highly variable genes. Cell clustering was determined using the “FindNeighbors” and “FindCluster” functions in the Seurat package. To investigate the proliferation of genes involved in the development of muscle stem cells, the “Monocle2” package was applied for conducting pseudotime analysis. The biological function analysis was performed using GO analysis. The scRNA-seq data have been deposited under accession number GSE250228.

### Quantification and statistical analysis

All data are represented as the mean of biological replicates, with individual sample data shown as dots. Values are reported as mean ± standard error of the mean (SEM). Statistical analyses were performed using unpaired Student’s t-tests or non-repeated measures analysis of variance (ANOVA), followed by Tukey’s test, using GraphPad Prism (GraphPad Software, Inc., San Diego, CA, USA). P values for statistical significance are indicated as * P < 0.05, ** P < 0.01, *** P < 0.001, not detected (n.d.), or not significant (n.s.).

## Results

### Dusp13 is directly regulated by MyoD activity

We previously generated MyoD-KI mice that allow us to assess MYOD expression through tdTomato fluorescence intensity [23]. To identify new regulators directly controlled by MYOD activity, we utilized RNA-seq data from MyoD-low and MyoD-high expressing MuSCs from the MyoD-KI mice, as well as freshly isolated quiescent MuSCs from Pax7^YFP/YFP^ (Pax7-YFP) mice (Figure 1A). Initially, we visualized the differentially expressed genes (FDR < 0.05) between the MyoD-low and MyoD-high populations using a heatmap (Figure 1A). We found that 966 genes were significantly upregulated as *MyoD* levels increased (MyoD-KI Cluster2). To further narrow down the targets, we consulted a gene expression database of proliferating and differentiating MuSCs from *MyoD* KO mice [40]. We identified 1797 genes that were not upregulated in *MyoD* KO mice (MKO Cluster2 in Figure 1B). Through a Venn diagram comparison, we combined 505 genes from the MyoD-KI and MKO clusters. Additionally, when we included *MyoD* ChIP-seq data from ChIP-Atlas, we found that 35 genes were enriched (Figure 1C) [41,42]. Finally, out of these 35 genes, six genes (*Ank1*, *Dmpk*, *Dusp13*, *Jsrp1*, *Rapsn*, and *Stac3*) were induced by *MyoD* overexpression in mouse embryonic fibroblasts (MEFs) [43] (Figure 1C and 1D; Figure S1A and S1B). We also confirmed that *Dusp13* mRNA was predominantly enriched in the quadriceps muscles compared to other tissues (Figure S2A). Due to the modulation of *Dusp13* expression by *MyoD levels* (Figure 1E–1H), we analyzed Dusp13 mRNA expression in sorted MyoD-low and -high MuSCs from MyoD-KI mice by RNAscope fluorescent in situ hybridization. Dusp13 RNA puncta were not obviously visible in MyoD-low, but accumulated in MyoD-high MuSCs (Figure S3A).

**Figure 1.**
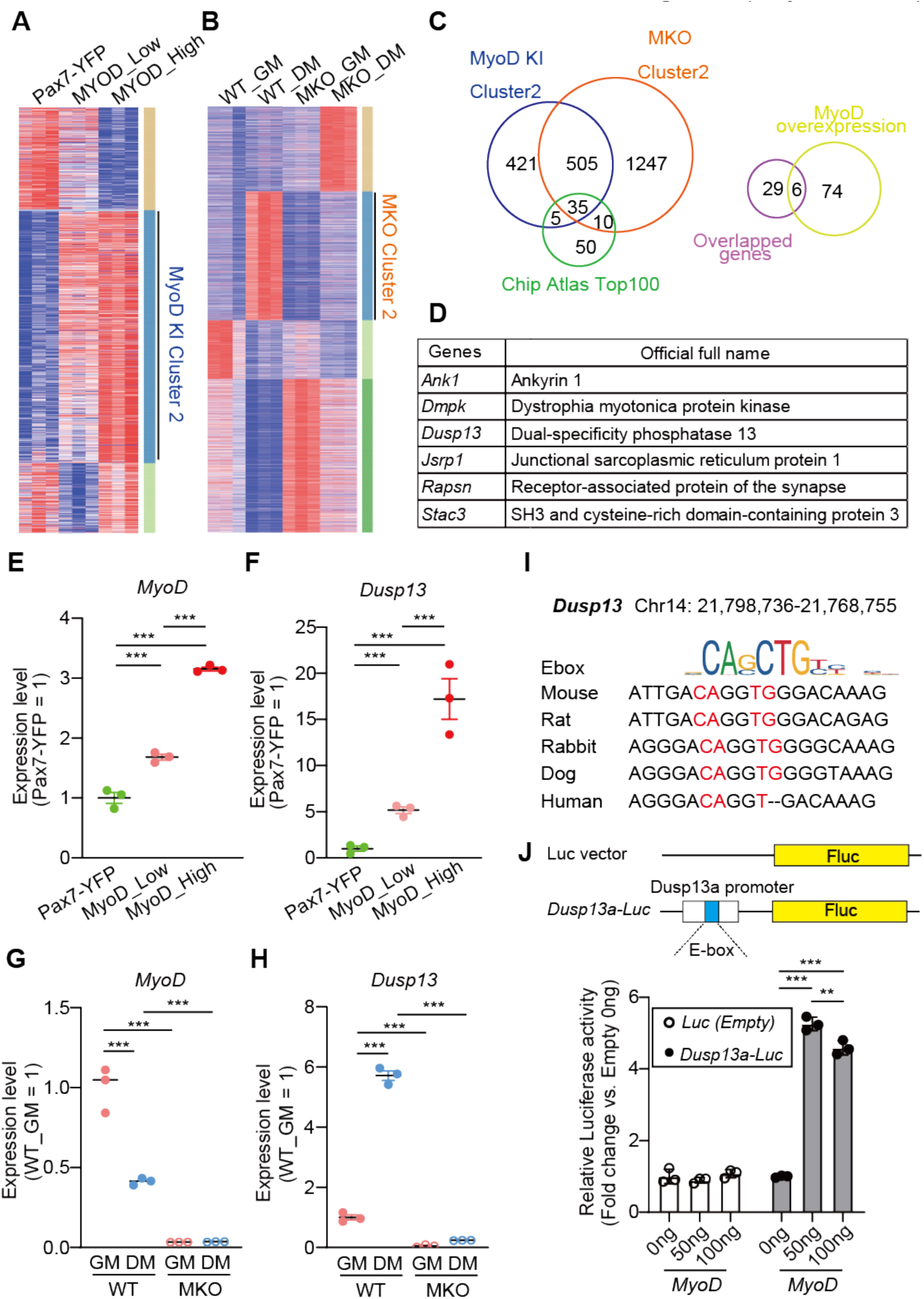
*Dusp13* is a novel target regulated by MyoD. **(A)** Clustering heatmap showing differentially expressed genes in RNA-seq data from Pax7-YFP (Quiescent MuSCs; QSCs), MyoD-low (MyoD-tdTomato low), and MyoD-high (MyoD-tdTomato high) (n = 3 per group). **(B)** Clustering heatmap of differentially expressed genes in RNA-seq between wild-type (WT) growth medium (GM) (WT_GM), *MyoD* knock-out GM (MKO_GM), WT differentiation medium (WT_DM), and MKO_DM (n = 3 per group). **(C)** Venn diagrams of indicated clusters from (**A, B)** and ChIP atlas top 100 genes potentially regulated by *MyoD* (Left). Venn diagrams of overlapped 35 genes (left) and 80 genes upregulated by *MyoD* overexpression in MEFs (Right). **(D)** List of six genes from (**C**). **(E, F)** Gene expression of *MyoD* (**E**) or *Dusp13* (**F**) analyzed by RNA-seq in the Pax7-YFP, MyoD-low, and MyoD-high MuSCs (n = 3 per group). **(G, H)** Gene expression of *MyoD* (**G**) or *Dusp13* (**H**) analyzed by RNA-seq in the WT_GM, WT_DM, MKO_GM, and MKO_DM groups (n = 3 per group). **(I)** E box-like sequences in the promoter region of *Dusp13* in mammalian species. **(J)** Luciferase reporter constructs of *Dusp13* upstream region (−150 to +50) containing an E box-like sequence (Top). Relative *Dusp13*-luciferase reporter activities in HEK293T cells overexpressing *MyoD* (Bottom) (n = 3 per group). All data are presented as mean ± standard error of the mean (SEM) P values calculated using edge R **(E, F)** (FDR-corrected), DEseq2 **(G, H)** (FDR-corrected), or Tukey’s test **(J)**; **P < 0.01, ***P < 0.001, n.s., not significant.

In addition, we identified a highly conserved E-box site 150 bp upstream of the transcription initiation site of *Dusp13*, according to the UCSC Genome Browser (Figure 1I). To determine if MYOD binds to the E-box of *Dusp13*, we ligated the putative E-box site of *Dusp13* to a pGL4.10 vector to generate a luciferase-*Dusp13*-E-box vector. We then co-transfected the plasmid with a *MyoD* expression vector into the HEK293T cell line. The luciferase activity was significantly upregulated in the presence of *MyoD* when the *Dusp13*-Ebox was present (Figure 1J). In contrast, the luciferase activity was not upregulated in the presence of *Pax7* or *Myogenin* (Figure S2C and S2D). These results suggest that *Dusp13* is a new target of MYOD during myogenesis.

### Single Dusp13 or Dusp27 KO mice exhibits no abnormality in muscle development or regeneration

To investigate the role of DUSP13 in skeletal muscle, we generated *Dusp13* KO mice using CRISPR-Cas9 technology. We confirmed that *Dusp13* expression in skeletal muscle was depleted in these *Dusp13* KO mice (Figure S2E). The *Dusp13* KO mice were born according to Mendelian laws and grew normally, similar to the litter mate wild type (WT) control mice, with respect to body weight and muscle cross-sectional area (CSA) of the soleus (SOL) and plantaris (PLA) muscles (Figure 2A–2C).

**Figure 2.**
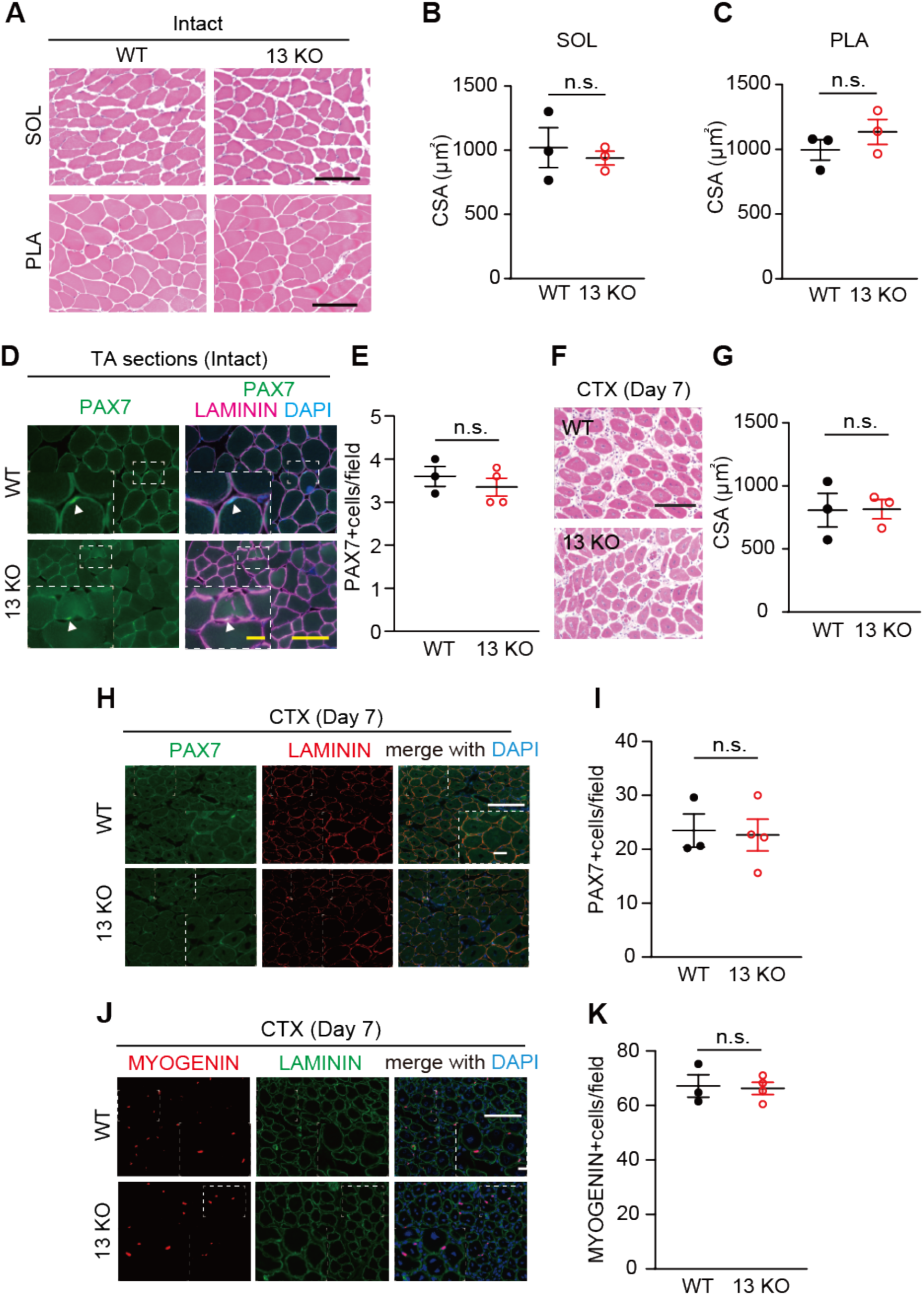
Absence of skeletal muscle phenotype in *Dusp13* knock-out (13 KO) mice during development and regeneration. **(A)** Hematoxylin and eosin (H&E) staining of Soleus (SOL) and Plantaris (PLA) muscle cross-sections of male *Dusp13* knock-out (13 KO) and WT mice at 10 weeks. Scale bar: 100 μm. **(B, C)** Cross-sectional area (CSA) of SOL (B) or PLA (C) (n = 3 per group). **(D)** Immunofluorescence of PAX7 and LAMININ in intact TA muscles. Scale bar: 100 µm. The area in the white dotted boxes is shown at a higher magnification. Scale bar: 25 μm. **(E)** Average number of PAX7 (+) nuclei in DAPI (+) in **(D)**. **(F)** H&E staining of tibialis anterior (TA) muscle cross-sections of male 13 KO and WT mice at 12 weeks, 7 days post cardiotoxin (CTX) injury. Scale bar: 100 µm. **(G)** CSA of damaged TA muscles (n = 3 per group). **(H)** Immunofluorescence of PAX7 with LAMININ on sections of TA muscles from male WT and *Dusp13* KO mice, 7 days post-CTX injury. Nuclei are stained with DAPI. Scale bar: 100 µm. The area in the white dotted boxes is shown at a higher magnification. Scale bar: 25 μm. **(I)** Average number of PAX7 (+) nuclei in **(H)** are quantified (n = 3–4 independent experiments). **(J)** Immunofluorescence of MYOGENIN with LAMININ on sections of TA muscle from male WT and 13 KO mice, 7 days post-CTX injury. Nuclei are stained with DAPI. Scale bar: 100 µm. The area in the white dotted boxes is shown at a higher magnification. Scale bar: 25 μm. **(K)** Average number of MYOGENIN (+) nuclei in **(J)** are quantified (n = 3–4 independent experiments). All data are presented as mean ± standard error of the mean (SEM) P values calculated using Student’s t-test; n.s., not significant.

The number of PAX7+ MuSCs in *Dusp13* KO mice is identical with WT mice (Figure 2D and 2E). We then investigated the regenerative ability in the *Dusp13* KO mice. We injected cardiotoxin (CTX) into the tibialis anterior (TA) muscles of both WT and *Dusp13* KO mice and no significant histological differences were observed in the CSA of the damaged myofibers with central nuclei at day 7 after the injury (Figure 2F and 2G). Furthermore, immunohistochemical analysis for PAX7 and MYOGENIN showed that the number of PAX7 or MYOGENIN (+) nuclei was similar between WT and *Dusp13* KO mice (Figure 2H–2K). The findings indicate that there is no significant impairment of muscle regeneration in the absence of *Dusp13*.

We hypothesized that other members of the *Dusp* family expressed in skeletal muscles could compensate for the function of DUSP13 upon *Dusp13* deletion. Within overlapping genes between the MyoD-KI Cluster 2 and MKO Cluster 2 in Figure 1A and 1B, we identified *Dusp27* as another member of the *Dusp* family that is expressed abundantly in the quadriceps muscles and the heart (Figure S2B). *Dusp27* expression was significantly higher in *Dusp13* KO muscles compared to WT muscles, suggesting compensatory induction following the loss of *Dusp13* (Figure 3A). We also observed a significant increase in *Dusp27* expression when *MyoD* expression increased (Figure 3B, Figure S3B), similar to *Dusp13* (Figure 1F, Figure S3A). Interestingly, we found a potential E-box site located 10 bp upstream of the transcription initiation site of *Dusp27*, which is highly conserved among multiple mammalian species (Figure 3C). The luciferase activity of luciferase-Dusp27-E-box was upregulated significantly in the presence of *MyoD* (Figure 3D) but not in the presence of *Pax7* or *Myogenin* (Figure S2C and S2D), suggesting that both *Dusp13* and *Dusp27* are regulated directly by MYOD activity. To test our hypothesis that DUSP27 compensates for the loss of *Dusp13*, we first generated *Dusp27* KO mice using CRISPR-Cas9. We confirmed the deletion of the gene through qPCR analysis (Figure S2F). Similar to the *Dusp13* KO mice, the *Dusp27* KO mice did not show any differences in body weight (data not shown), myofiber CSA in intact muscles (Figure 3E-H), number of PAX7+ MuSCs in intact muscles (Figure 3I and 3J), myofiber CSA after muscle CTX injury (Figure 3K and 3L), and the number of myogenic populations labeled by either PAX7 or MYOGENIN after CTX injury (Figure 3M–3P). These results demonstrate that neither *Dusp13* nor *Dusp27* deficiency influences muscle development and acute muscle regeneration.

**Figure 3.**
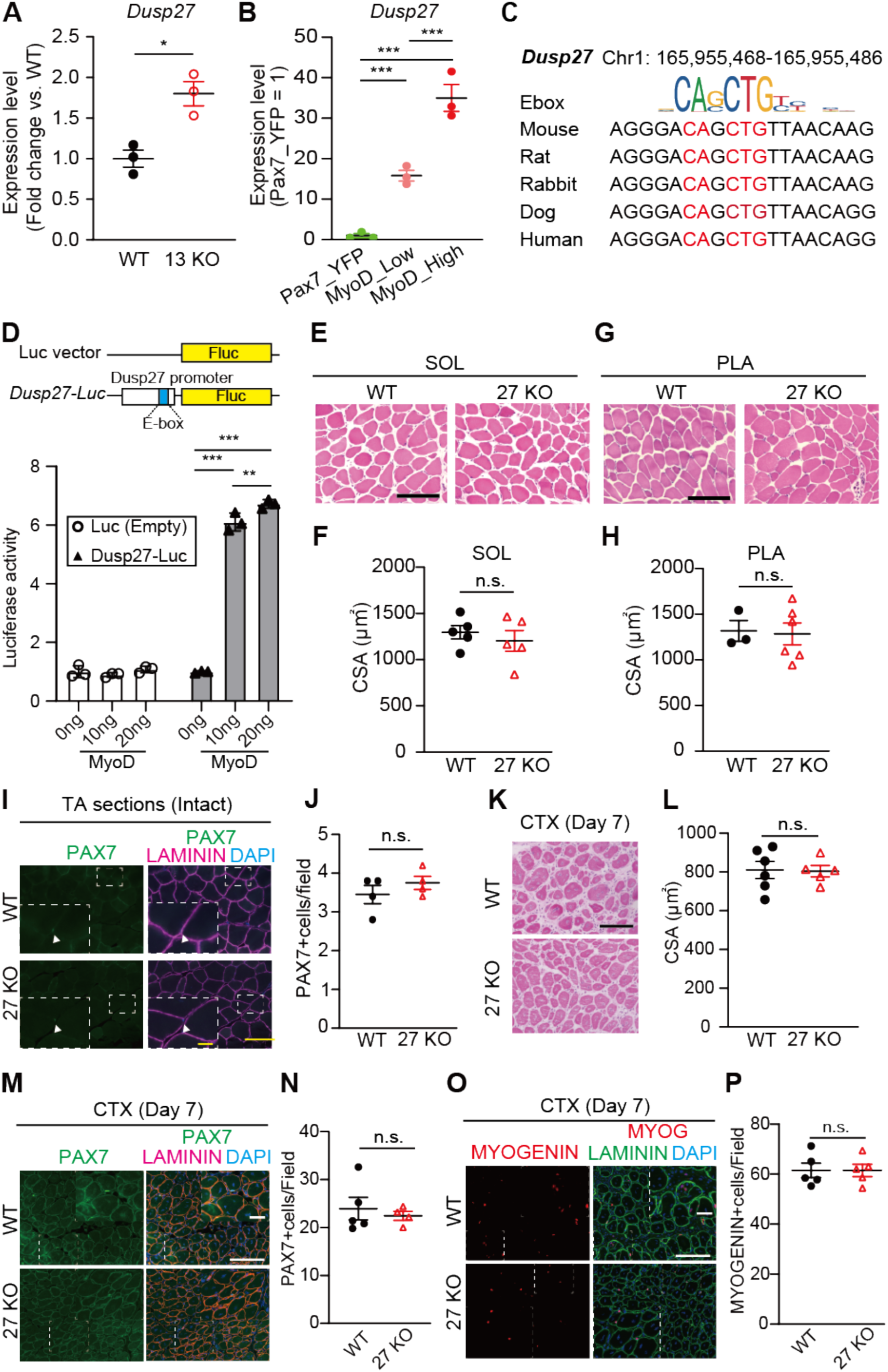
*Dusp27* KO mice exhibit comparable muscle development and regeneration to WT mice. **(A)** qPCR analysis of *Dusp27* mRNA levels in gastrocnemius muscles of male WT and 13 KO mice at 12 weeks (n = 3 per group). **(B)** Gene expression of *Dusp27* in RNA-seq data from Pax7-YFP, MYOD-low, and MYOD-high groups (n = 3 per group). **(C)** Ebox-like sequences in the promoter region of *Dusp27* in mammalian species. **(D)** Luciferase reporter constructs of *Dusp27* upstream region (−1,000 to +200) containing an Ebox-like sequence (Top). Relative *Dusp27*-luciferase reporter activities in HEK293T cells overexpressing MyoD (Bottom) (n = 3 per group). **(E, G)** H&E staining of SOL and PLA muscle cross-sections. Scale bar: 100 μm. **(F, H)** CSA of SOL **(F)** or PLA **(H)** muscles of male *Dusp27* KO (27 KO) and WT (n = 3–6 independent experiments) mice at 12 weeks. **(I)** Immunofluorescence of PAX7 and LAMININ in intact TA muscles. Scale bar: 100 µm. The area in the white dotted boxes is shown at a higher magnification. Scale bar: 25 μm. **(J)** Average number of PAX7 (+) nuclei in DAPI (+) in **(I)**. **(K)** H&E staining of TA muscle cross-sections of male 27 KO and WT mice at 12 weeks (n = 5–6 independent experiments), 7 days post-cardiotoxin injury. Scale bar: 100 µm. **(L)** CSA of damaged TA muscles. **(M, N)** Immunofluorescence of PAX7 **(M)** or MYOGENIN **(N)** with LAMININ on sections of TA muscles from male WT and 27 KO mice, 7 days post-CTX injury. Nuclei are stained with DAPI. Scale bar: 100 µm. The area in the white dotted boxes is shown at a higher magnification. Scale bar: 25 μm. **(O, P)** Average number of PAX7 (+) or MYOGENIN (+) nuclei in **(O)** and **(P)** are quantified (n = 5– 8 independent experiments). All data are presented as mean ± standard error of the mean (SEM) P values calculated using Student’s t-test, Tukey’s test, and edge R; *P < 0.05, **P < 0.01, ***P < 0.001, n.s., not significant.

### Muscle regeneration is delayed in Dusp13:Dusp27 DKO mice

To investigate the role of DUSP13 and DUSP27 in skeletal muscles, we generated DKO mice. The body weights and muscle weights of DKO mice were found to be similar to those of age-matched WT mice (Figure S4A and S4B). Furthermore, both the SOL and PLA muscles exhibited normal myofiber size (Figure S4C–S4E), and the number of MuSCs (Figure S4F-I) in DKO mice was also normal, as in WT mice. However, histological analysis on days 3, 7, and 28 after CTX injury clearly demonstrated the essential role of DUSP13 and DUSP27 in muscle regeneration (Figure 4A–4D). Specifically, seven days post-injury, we observed a notable increase in PAX7 (+) cells and a decrease in MYOGENIN (+) cells (Figure 4E, 4F, 4H, and 4I) in DKO muscles compared to those in WT mice. Furthermore, the percentage of Ki67 (+) cells in MYOGENIN (+) cells was increased significantly in DKO muscles (Figure 4G and 4J). At 28 days post-injury, when MuSCs are returning to their niche after completing the muscle regeneration process, we observed a trend towards a reduction in the number of PAX7 (+) cells (Figure S4J and S4K), accompanied by a notable increase in the number of perduring MYOGENIN (+) cells undergoing regeneration (Figure S4L and S4M).

**Figure 4.**
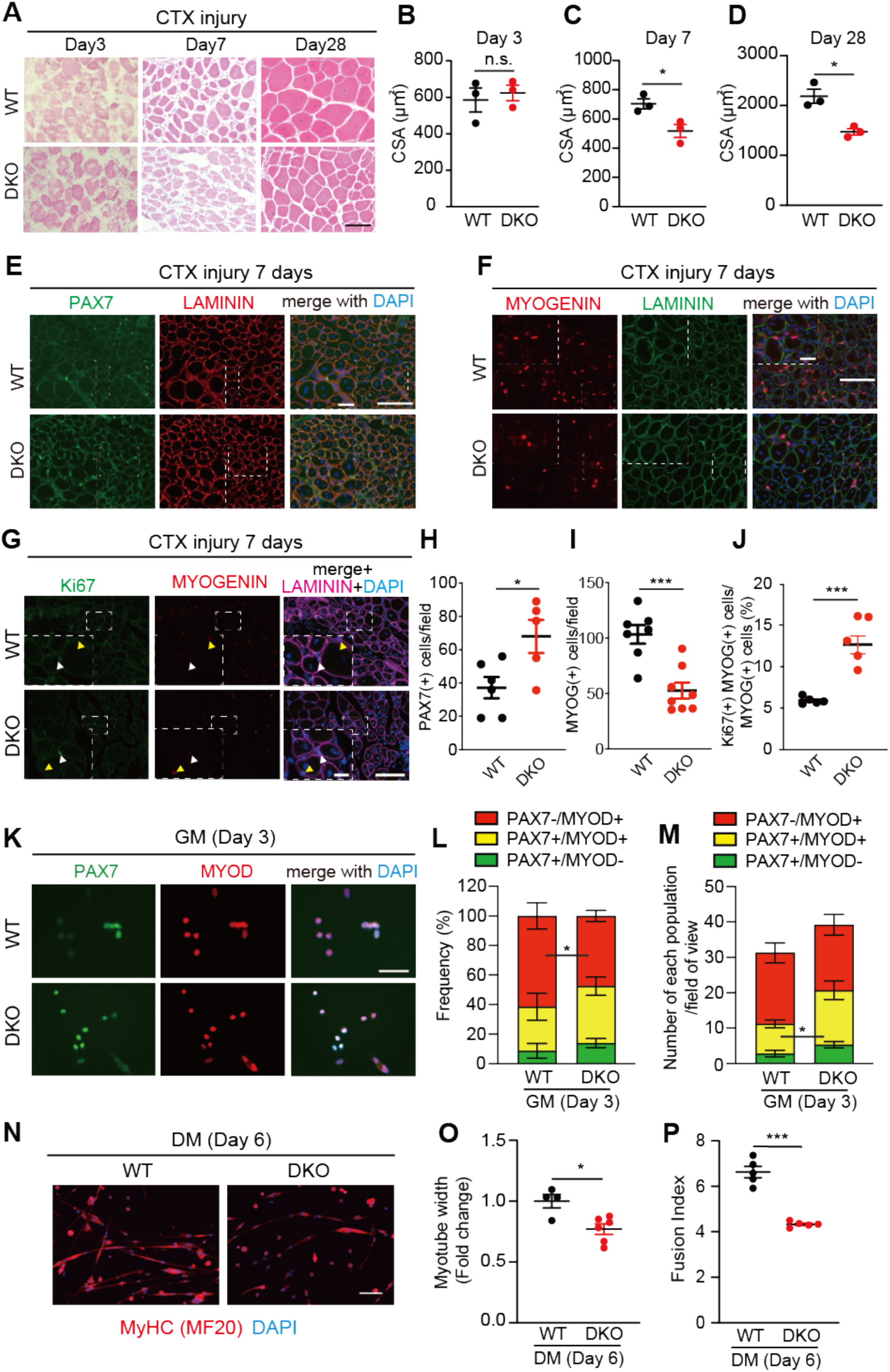
Loss of both *Dusp13* and *Dusp27* impairs muscle regeneration after acute muscle injury. **(A)** Hematoxylin and eosin (H&E) staining of tibialis anterior (TA) muscle cross-sections of male *Dusp13* and *Dusp27* double knock-out (DKO) (n = 3) and WT (n = 3) mice at 12–15 weeks of age, 3, 7, and 28 days post**-**cardiotoxin (CTX) injury. Scale bar: 100 µm. **(B–D)** Cross-sectional area (CSA) of damaged TA muscles at indicated time points. **(E, F)** Immunofluorescence of PAX7 (**E**) or MYOGENIN (**F**) with LAMININ on sections of TA muscles from male WT and DKO mice, 7 days post**-**CTX injury. Nuclei are stained with DAPI. Scale bar: 100 µm. The area in white dotted boxes is shown at a higher magnification. Scale bar: 25 μm. **(G)** Iimmunofluorescence of KI67 and MYOGENIN with LAMININ on sections of TA muscles from male WT and DKO mice, 7 days post**-**CTX injury. Nuclei are stained with DAPI. Scale bar: 100 µm. The area in white dotted boxes is shown at a higher magnification. Scale bar: 25 μm. White arrowhead indicates KI67+/MYOGENIN+ cells. Yellow arrowhead indicates KI67-/MYOGENIN+ cells. **(H, I)** Average number of PAX7 (+) or MYOGENIN (+) nuclei in **(E)** and **(F)** are quantified (n = 5– 8 independent experiments). **(J)** Frequency of Ki67 (+) nuclei in MYOGENIN (+) in **(G)** are quantified (n = 5 per group) **(K)** Immunofluorescence of PAX7 and MYOD on cultured MuSCs from female WT and DKO mice. Nuclei are stained with DAPI. Scale bar: 100 µm. **(L, M)** Quantification of the proportion **(L)** and numbers **(M)** of MuSCs undergoing self-renewal (PAX7 (+), MYOD (-)), activation (PAX7 (+), MYOD (+)), and differentiation [PAX7 (+), MYOD (-)] in (**K**). (n = 6 independent experiments) **(N)** Immunofluorescence with an antibody against myosin heavy chain (MyHC) on MuSCs differentiated for six days in DM. Nuclei are stained with DAPI. Scale bar: 100 µm. **(O, P)** Myotube width **(O)** and Fusion index (numbers of myonuclei per MyHC (+) myotube) **(P)** in **(N)** are quantified (50 myotubes analyzed per experiment). All data are presented as mean ± standard error of the mean (SEM). P values calculated using Student’s t-test; *P < 0.05, **P < 0.01, ***P < 0.001, n.s., not significant.

To confirm our *in vivo* findings indicating a severe delay in muscle regeneration in DKO mice, we isolated MuSCs from DKO and WT mice using magnetic cell sorting (MACS) with α7-integrin microbeads [6,23] and cultured them for three days. Immunofluorescent analysis using antibodies against PAX7 and MYOD showed a lower proportion of PAX7 (-) and MYOD (+) cells and an increased number of PAX7 (+) and MYOD (+) cells in the DKO mice (Figure 4K–4M), indicating that DUSP13/DUSP27 are necessary for the transition from proliferation to differentiation during myogenesis. In addition, we observed a higher number of proliferating Ki67 (+) MuSCs in DKO mice (Figure S5A–S5C). In addition, both myotube width and fusion index exhibited a significant reduction in DKO mice relative to in WT mice after six days of MuSC differentiation (Figure 4N–4P). The findings suggest that DUSP13 and DUSP27 dictate the balance between the proliferation and differentiation of MuSCs during myogenesis.

### Single-cell RNA sequencing revealed that Dusp13 and Dusp27 are highly expressed in the late differentiation stage of MuSCs

While MYOD activates *Dusp13* and *Dusp27* through their E-box, and *DKO* mice exhibit diminished muscle regeneration activity, the molecular mechanisms by which DUSP13 and DUSP27 regulate myogenesis remain elusive. Additionally, the expression patterns of *Dusp13* and *Dusp27* in different stages of myogenic progenitors within the highly heterogeneous MuSC populations are not well-defined. To gain a more comprehensive understanding of the molecular signatures of *Dusp13* and *Dusp27*-expressing myogenic cells, we conducted single-cell RNA sequencing (scRNA-seq) on MyoD-high expressing cells isolated from cultured MuSCs obtained from MyoD-KI mice (Figure 5A). The approach allowed us to enrich activated/differentiated MyoD (+) cells [23]. MuSCs from MyoD-KI mice were cultured for five days in a growth medium, and only live MyoD-high MuSCs with PI (-) negative populations were sorted by fluorescence-activated cell sorter (FACS) for subsequent scRNA-seq library preparation (Figure 5A). Utilizing the Seurat single-cell analysis package, we identified five distinct clusters of cells (C0–C4) presented in the uniform manifold approximation and projection (UMAP) (Figure 5B). Each cluster exhibited a unique transcriptomic fingerprint characterized by the top 10 differentially expressed genes (Figure S6A).

**Figure 5.**
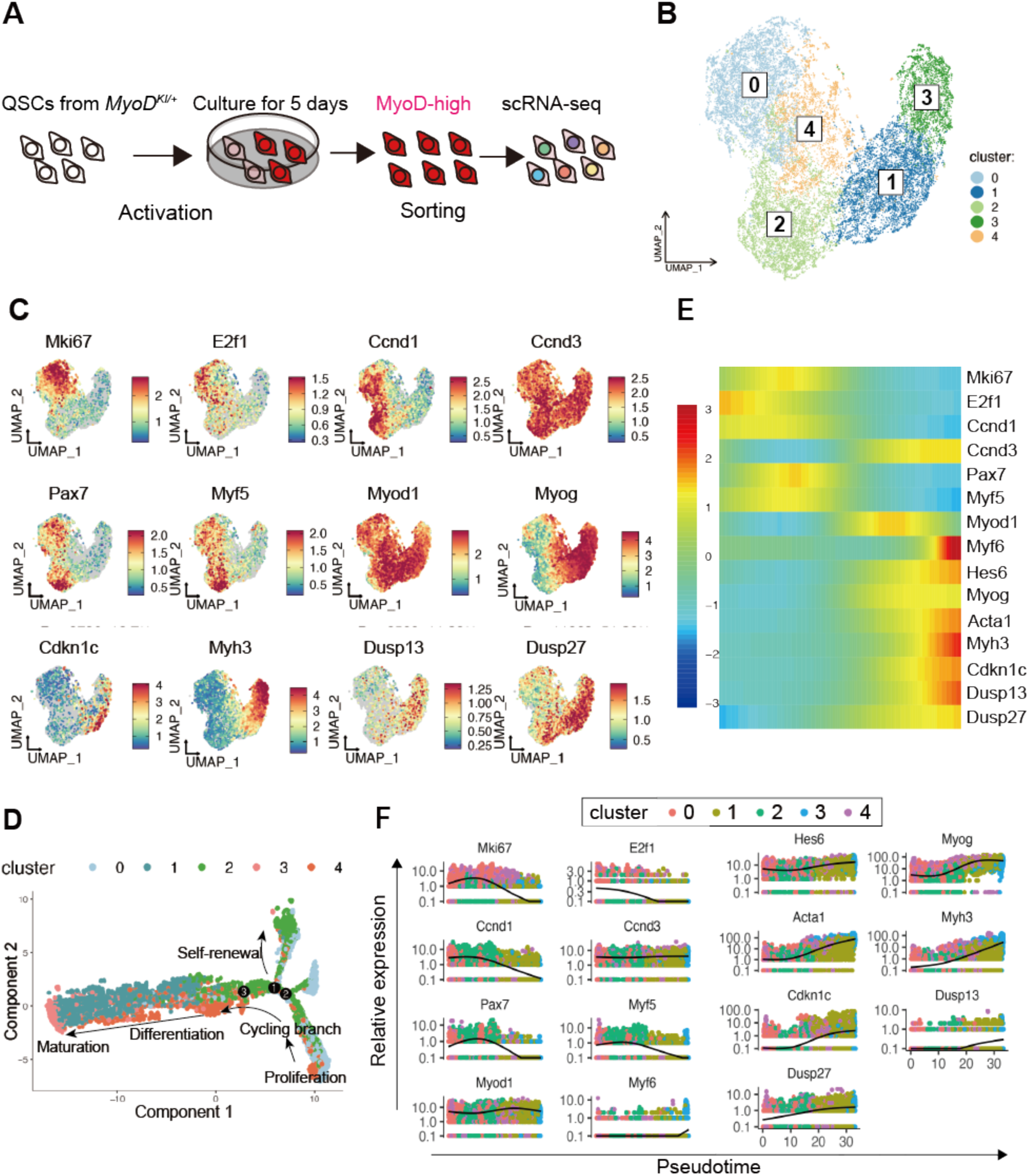
Single-cell RNA-sequencing analysis of MyoD-high MuSCs from MyoD-KI mice reveals *Dusp13* and *Dusp27* expression in the late stage of MuSCs. **(A)** Schematic diagram illustrating the scRNA-seq analysis of MyoD-high MuSCs from MyoD-KI mice. **(B)** Transcriptomic atlas and clustering of the subset of MyoD-high MuSCs from MyoD-KI mice. **(C)** Single-cell expression pattern of the target genes. **(D)** Pseudotime analysis performed using the Monocle package. The plot depicts the hierarchy of MyoD-high MuSCs in the pseudotime trajectory. **(E)** A heatmap showing the expression of representative genes in the pseudotime analysis. Each column represents a cell, and each row represents a gene. **(F)** Pseudotime-ordered single-cell expression of representative genes.

To further understand the characteristics of the clusters, a Gene Ontology (GO) analysis on differentially expressed genes (DEGs) was conducted among the clusters (Figure S6B–S6F). C0 and C4 showed upregulation of genes associated with chromosome segregation and DNA replication (Figure S6B and S6F). C1 and C3 displayed strong enrichment for terms related to muscle differentiation, maturation, and contraction (Figure S6C and S6E). C2 was characterized by genes associated with ameboidal-type cell migration and extracellular organization (Figure S6D), akin to the previously reported MyoD-low MuSC signatures [23]. Next, we investigated cell cycle and myogenic markers, including *MKi67, E2f1, Ccnd1, Ccnd3, Pax7, Myf5, Myod1, Myogenin, Cdkn1c (p21), Myh3, Dusp13*, *and Dusup27* on UMAP (Figure 5C). *MyoD* is expressed ubiquitously across all clusters but with variations between the clusters (Figure 5C). *Mki67* and *E2f1*, which are proliferative markers, were highly expressed in C0 and C4 but not in C1, C2, and C3 (Figure 5C). Undifferentiated MuSC markers, *Pax7* and *Myf5*, were enriched exclusively in C0 and C2 (Figure 5C). Notably, *Dusp13* and *Dusp27* exhibited pronounced enrichment in C1 and C3, characterized by upregulation of genes associated with myogenic differentiation and maturation (Figure 5C). Considering the GO term and UMAP, C0 and C4 represent proliferative states, while C1 and C3 contain the differentiation and maturation state of MuSCs. Conversely, C2 reflects the self-renewing MuSCs returning to the undifferentiated state.

Furthermore, we used Monocle to analyze the MYOD-high expressing MuSC clusters and infer MuSC state trajectories (Figure 5D–5F). The cell state trajectory model revealed three branches for the MyoD-high-expressing MuSCs. The branches suggested that some clusters were transitioning toward further proliferation (C1, C4), myogenic differentiation/maturation (C1, C3), or self-renewal (C2) to return to an undifferentiated quiescent state (Figure 5D). Subsequently, we focused the Monocle analysis on selected genes related to myogenic development/regeneration and cell cycle (*Pax7, Myf5, Myod1, Myogenin, Myf6, Myh3, Myh4, Hes6, Acta1, ccnd1, ccnd3, E2f1, Mki67, Cdkn1c [p21], Dusp13, and Dusp27*). Cluster 3 contained *Dusp13* and *Dusp27*, along with *Myogenin, Myh3, Hes6, Acta1,* and *Cdkn1c (p21)* (Figure 5E and 5F). When visualizing the pseudotime progression axis of these selected genes, we observed that cells from C0 and C4, which expressed genes associated with the cell cycle (*Mki67* and *E2f1*) but had lower expression of *Pax7* and *Myf5*, were located at the initiation point of pseudotime (Figure 5E and 5F). Cells from C2, which showed lower levels of *Mki67* and *E2f1* but higher levels of *Pax7*, *Myf5*, and *Ccnd1*, were found mainly in the middle point of pseudotime, suggesting that C2 represents a self-renewing MuSCs state [23,44]. In contrast, cells from C1 and C3, which highly expressed genes associated with myogenic differentiation and maturation, were located distally at the endpoint of pseudotime (Figure 5E and 5F), indicating myogenic differentiation and maturation states (Figure 5E and 5F). *Dusp13* and *Dusp27*, enriched in C1 and C3, progressed alongside myogenic differentiation markers and *Cdkn1c (p21)* (Figure 5E and 5F). To further support our scRNA-seq data, we visualized *Dusp13* and *Dusp27* RNA in different stages of myogenic cells (freshly isolate, GM6, and DM6) using RNAscope (Figure S7A and S7B). The results revealed that puncta for both genes were either not visible or present in low numbers in the freshly isolated PAX7+ MuSCs but increased gradually as myogenesis progressed (Figure S7A and S7B). Overall, clustering and cell state trajectory analysis based on scRNA-seq data of MyoD-high-expressing MuSCs and the RNAscope analysis suggest that *Dusp13* and *Dusp27* are enriched only in the late stages of MyoD-positive MuSCs co-expressing *Myogenin*, *Myh3*, and *Cdkn1c*. This indicates that these genes may regulate the transition from a proliferative state to a myogenic differentiation state.

### Dusp13 drives precocious muscle differentiation *in MuSCs*

Since our scRNA-seq analysis of MyoD-high expressing cells indicates that *Dusp13* and *Dusp27* are expressed in the subset of cells expressing late myogenic markers, such as myogenin, to investigate whether DUSP13 plays a role in initiating the differentiation process during myogenesis, we transduced FACS-isolated MuSCs from adult Pax7-YFP mice with an adenovirus vector expressing *Dusp13* or GFP under the CMV promoter (Figure 6A, Figure S8A, and S8B). Two days post-infection, we observed a significant reduction in the number of MuSCs transduced with *Dusp13* compared to the GFP control (Figure 6B). Notably, immunofluorescent analysis using antibodies against PAX7 and MYOGENIN showed that *Dusp13* overexpression led to induction of MYOGENIN expression in PAX7 (+) cells, which are typically not detected under normal culture conditions (Figure 6C and 6D). However, overexpression of *Dusp13* did not alter MyoD expression (Figure S8C and S8D). Furthermore, when we cultured MuSCs overexpressing *Dusp13* for seven days and assessed their myotube formation ability, we found that both the myotube width and fusion index were lower compared to the GFP control (Figure 6E–6H). These results suggest that overexpression of *Dusp13* in activated MuSCs induces precocious differentiation, which inhibits the amplification of MuSCs and results in smaller myotube size and lower fusion index.

**Figure 6.**
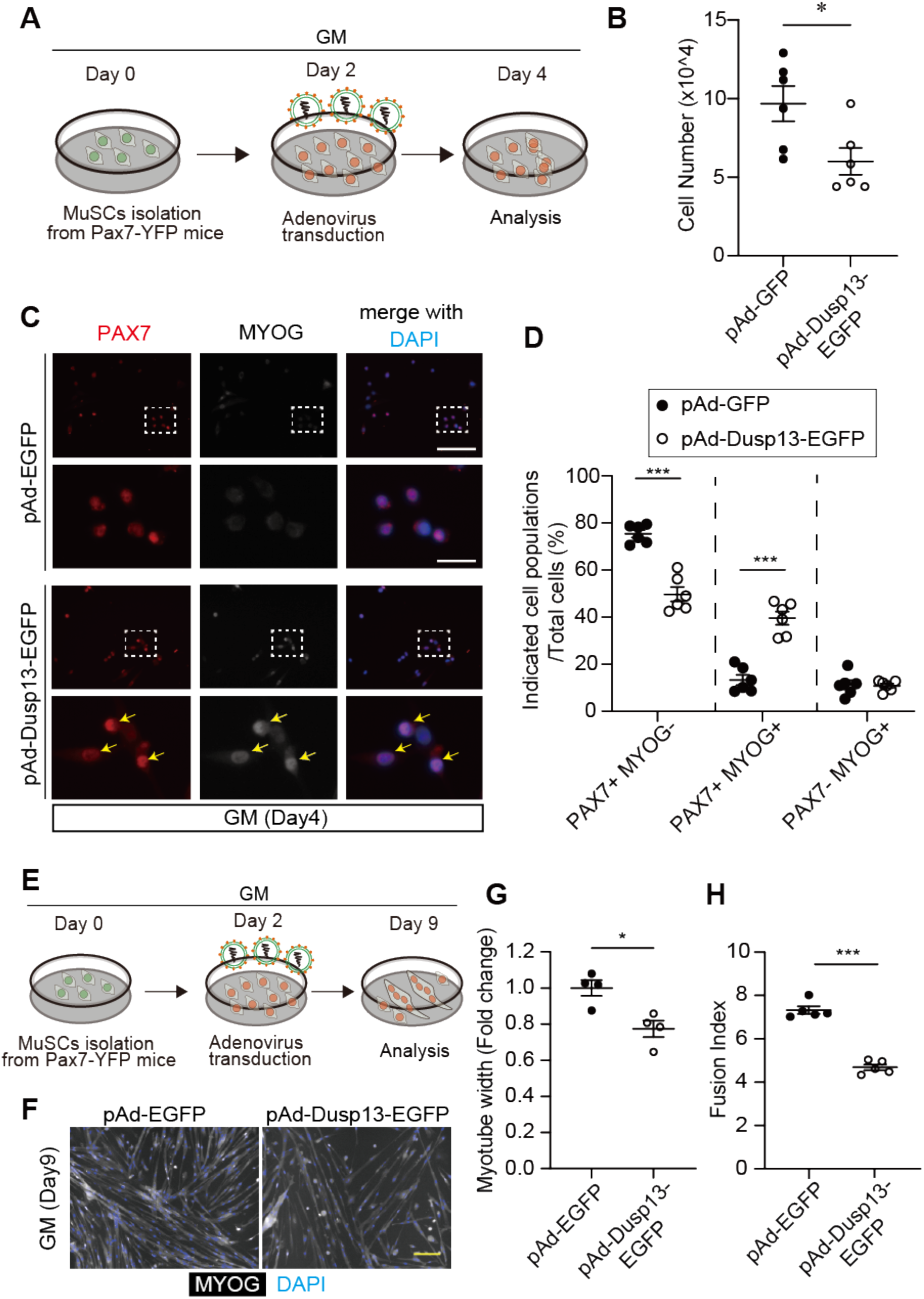
Overexpression of *Dusp13* induces precocious muscle differentiation independent of phosphatase activity. **(A)** Schematic diagram illustrating adenovirus infection into MuSCs from 15–16-week-old male Pax7-YFP mice on day 4 of growth medium (GM) culture. **(B)** Number of cultured MuSCs after transduction with adenovirus vectors overexpressing GFP (pAd-EGFP) or Dusp13 (pAd-Dusp13-EGFP). **(C)** Immunofluorescence staining of PAX7 and MYOGENIN in cultured MuSCs after transduction with adenovirus vectors overexpressing GFP (pAd-EGFP) or Dusp13 (pAd-Dusp13-EGFP) on GM day 4. Nuclei are stained with DAPI. The yellow arrow indicates PAX7 (+)/MYOGENIN (+) cells. (n = 6 per group). Scale bar: 100 µm (Upper); 25 µm (Bottom). **(D)** The proportion of each population in (**C**). **(E)** Schematic diagram illustrating adenovirus infection into MuSCs from 15–16-week-old male Pax7-YFP mice on GM day 9. **(F)** Immunofluorescence staining of MYOGENIN in cultured MuSCs after transduction with adenovirus vectors overexpressing GFP (pAd-EGFP) or Dusp13 (pAd-Dusp13-EGFP) on GM day 9. Nuclei are stained with DAPI. Scale bar: 100 µm **(G, H)** Myotube width **(G)** and Fusion index (numbers of myonuclei per myotube) **(H)** in **(F)** are quantified (50 myotubes analyzed per each experiment) All data are presented as mean ± standard error of the mean (SEM). P values calculated using Student’s t-test; *P < 0.05, **P < 0.01, ***P < 0.001, n.s., not significant.

To investigate whether the phosphatase activity of DUSP13 is involved in myogenic induction, we transduced MuSCs from adult PAX7-YFP mice with pAd-Dusp13_Mut, which lacks enzymatic activity. Surprisingly, we discovered that DUSP13 without phosphatase activity still enhanced the population of PAX7 (+) and MYOGENIN (+) cells to the same extent as normal DUSP13 (Figure S8E and S8F). Overall, the results suggest that DUSP13 acts as a driver for myogenin during myogenic progression, which is mediated by phosphatase activity-independent mechanisms.

## Discussion

Skeletal muscle regeneration relies heavily on the activity of muscle stem cells (MuSCs). The regulation of MuSC dynamics is mediated by the sequential expression of various transcription factors during myogenesis. MYOD is essential for myogenic lineage specification during embryogenesis [12,45,46]. In adult mice regeneration, MYOD is induced in activated MuSCs, modulating the balance between self-renewal and differentiation [21,22,47–49]. However, the molecular mechanisms by which MYOD governs fate determination in MuSCs remain unclear, particularly regarding its direct targets beyond *Myogenin*, which have not been fully identified. Therefore, it is crucial to identify additional direct targets of MYOD that play key roles in skeletal myogenesis during development and regeneration. The diverse and highly heterogeneous nature of MuSCs, particularly during myogenesis, has posed a challenge in deciphering molecular signatures in MYOD-expressing MuSCs due to a lack of appropriate tools. This has hindered the elucidation of MYOD’s role in fate determination. To address this, we previously engineered MyoD-KI mice, providing a platform to distinguish and isolate undifferentiated (MyoD-low) and committed MuSCs (MyoD-high).

In the present study, we integrated our previously established whole-transcriptome data encompassing quiescent, undifferentiated (MyoD-low), and committed (MyoD-high) MuSCs from MyoD-KI reporter mice [23] with publicly available databases from *MyoD* KO mice [40], Chip-atlas [41,42], and *MyoD* overexpression [43]. Through this integration, we identified *Dusp13* as a novel molecular switch regulated by MyoD, stimulating the initiation of myogenic differentiation. To understand the function of DUSP13, we generated *Dusp13* KO mice using CRISPR-Cas9. Surprisingly, muscle development and regeneration abilities in *Dusp13* KO mice mirrored those of their WT counterparts. This led us to investigate the compensatory role of other members of the DUSP family. We found that *Dusp27* was also directly regulated by MyoD. Although *Dusp27* KO mice generated in the present study did not exhibit marked differences in muscle development and regenerative ability following CTX injury, the combined deficiency of *Dusp13* and *Dusp27* in mice resulted in substantial delays in muscle regeneration after CTX injury. The findings suggest a compensatory interplay between DUSP13 and DUSP27 during muscle regeneration. The quantities of PAX7 (+) quiescent MuSCs were identical between WT and DKO mice before injury. However, at 28 days post-CTX injury, when muscle regeneration was almost complete, DKO mice had lower levels of PAX7 (+) cells compared to WT mice. Conversely, DKO mice had more MYOGENIN (+) cells remaining at day 28 post-injury compared to WT mice. At day 7 post-injury, DKO mice had higher numbers of PAX7 (+) cells and fewer MYOGENIN (+) cells. The results suggest that the absence of *Dusp13* and *Dusp27* promotes the imbalance of the fate determination processes in MuSCs following muscle injury. *In vitro* analyses employing isolated MuSCs from DKO mice supported these findings, showing that the absence of *Dusp13* and *Dusp27* promotes proliferation rather than differentiation. Consequently, the delay/failure in transitioning from the proliferation phases to differentiation, attributed to the deficiency of *Dusp13* and *Dusp27*, disrupts the muscle repair process in adult mice.

To further investigate the regulatory dynamics of *Dusp13* and *Dusp27* in fate determination, we conducted scRNA-seq on MyoD-high expressing MuSCs isolated from MyoD-KI mice. Intriguingly, both *Dusp13* and *Dusp27* expression were enriched in specific clusters (C1 and C3) within the MyoD-high expressing MuSC populations. The clusters were characterized by increased expression of late myogenic markers, such as *Myogenin, Myh3,* and *Cdkn1a*, and decreased expression of *Pax7*, *MKi67*, and *Myf5*. Pseudotime cell state trajectory analysis was used to distinguish the fate determination of MuSCs during myogenesis. Cluster C0 represented a proliferative state with high expression of undifferentiated markers (*Pax7* and *Myf5*) and cell cycle markers (*MKi67* and *E2f1*), which are projected to further proliferate (C4), differentiate/mature (C1 and C3), or self-renew to return to an undifferentiated quiescent state (C2). We believe that the scRNA-seq data from our MyoD-KI mice will provide a valuable resource for uncovering novel molecular mechanisms involved in myogenic fate transition (proliferation, differentiation, or self-renewal/quiescence) of activated MuSCs during myogenesis.

Considering the limited expression of *Dusp13* and *Dusp27* in C1 and C3 (diifferentiate/mature clusters), we hypothesized that the two genes play a crucial role in the transition from proliferation to differentiation. In our study, we overexpressed *Dusp13* in activated MuSCs *in vitro* using an adenovirus vector. Although the expression of PAX7 and MYOGENIN is typically mutually exclusive during myogenesis [50], a small subset of cells exhibits positivity for both PAX7 and MYOGENIN, representing an intermediate population initiating differentiation [51]. In our model, a limited number of PAX7 (+) and MYOGENIN (+) cells were also identified. Notably, *Dusp13* overexpression augmented the populations of PAX7 (+) and MYOGENIN (+) cells significantly. Moreover, the promoter activities of both Dusp13 and Dusp27 were not upregulated by the *Myogenin* expression vector. The results collectively corroborate the findings of our scRNA-seq analysis, supporting the hypothesis that the MYOD-DUSP13/DUSP27-MYOGENIN regulatory networks also orchestrate the initiation of differentiation during muscle regeneration in adult mice.

There are limited studies on the regulatory roles of the DUSP family in myogenesis, leaving the molecular targets modulated by DUSP13 and DUSP27 in MuSCs currently unknown. The DUSPs family possesses the dual ability to dephosphorylate phosphor-serine/threonine and phospho-tryosoine residues. Currently, 25 different DUSPs are identified and categorized into six subfamilies [25]. Our single-cell RNA-seq analysis of MyoD-high MuSCs revealed enrichment of other DUSP family members such as *Dusp3* and *Dusp14* in C1 and C3 clusters (data not shown). While the relationship with Dusp13/27 was not investigated in this study, it is noteworthy that the expressions of *Dusp3* and *Dusp14* were not compensatory upregulated by the loss of *Dusp13* and *Dusp27*, suggesting that uncovering additional functions of other DUSP family members in myogenesis is warranted in future research.

DUSP13, an atypical DUSP, has versatile dephosphorylation substrates, including mitogen-activated protein kinases (MAPKs). Forced expression of pAd-Dusp13_Mut vector, which has no phosphatase activity, into MuSCs, led to the emergence of PAX7 (+) and MYOGENIN (+) cells *in vitro*, indicating phosphatase-independent mechanisms mediated by DUSP13 for myogenic differentiation. Interestingly, DUSP27 is recognized as a pseudo-phosphatase, characterized by catalytically inactive phosphatase activity. Various pseudo-phosphatases are reported to possess pivotal functions in different organisms and pathological conditions [33]. Inhibition of ERK1/2 promotes myogenic differentiation and fusion in both mice and avian models [52]. The DUSP family, lacking a kinetic-interacting motif, inhibits ERK1/2 signaling [53–55], suggesting that the phosphatase-independent activities of DUSP13 and DUSP27 potentially regulate ERK signaling in MuSCs. Future investigations are expected to unravel the detailed molecular mechanisms by which DUSP13 and DUSP27 orchestrate fate switching in activated MuSCs during myogenesis.

Although our study demonstrates that DKO mice exhibit reduced muscle regeneration capacity, several limitations must be considered when interpreting our results. The systemic deletion of *Dusp13* and *Dusp27* complicates the isolation of specific effects on muscle regeneration. To further investigate this, we isolated MuSCs from DKO mice and found impaired myotube formation compared to WT mice. However, optimal muscle regeneration requires the coordinated interaction of multiple cell types, including inflammatory cells [56,57] and fibro-adipogenic progenitors (FAPs) [58,59]. The expression of *Dusp13* and *Dusp27* is comparatively lower in other tissues than in skeletal muscles (Figure S2A– S2B). Furthermore, re-analysis of single-cell and -nucleus RNA-seq data from intact skeletal muscle indicates that *Dusp13* and *Dusp27* are highly enriched in myonuclei but not in other cell types, such as immune cells and endothelial cells [60]. Most FAPs are negative for *Dusp13* and *Dusp27*, although a small subset expresses the genes. Given the crucial role of FAPs in muscle regeneration [58,59,61], we cannot entirely exclude the possibility that the delayed muscle regeneration in DKO mice could partially result from the loss of *Dusp13* and *Dusp27* in the subset of FAPs. The area of infiltrated macrophages was not apparent between WT and DKO muscles post-injury (data not shown). Further investigations using *Pax7^CreERT2^* mice to specifically deplete *Dusp13* and *Dusp27* in MuSCs will be necessary to clarify this aspect. Another unresolved issue is why DKO mice exhibit no differences in skeletal muscle size, weight, or histology compared to WT mice under normal conditions. The lack of effect from the loss of DUSP13 and DUSP27 on muscle development during embryogenesis is unclear. It is possible that other DUSP family members compensate for their roles during embryonic and postnatal muscle formation, or that the downstream targets of DUSP13 and DUSP27 differ between muscle regeneration and embryonic myogenesis.

In summary, using our previously established MyoD-KI mice model, we identified *Dusp13* and *Dusp27* as novel MYOD target genes that play a pivotal role in the fate switching of MuSCs. Furthermore, we demonstrated that *Dusp13* and *Dusp27* expressions are highly enriched in differentiation clusters revealed by scRNA-seq of MyoD-high expressing MuSCs. Beyond *Dusp13* and *Dusp27* being novel determinants for the initiation of myogenic differentiation, further analysis of scRNA-seq data within each cluster would yield valuable insights into potential targets of DUSP13 or DUSP27 in MuSCs during myogenesis.

## Acknowledgments

We would like to thank Satoshi Yamazaki and Yuji Yamazaki from the Division of Stem Cell Therapy, University of Tsukuba, for their valuable technical assistance in cell sorting using flow cytometry. We also thank Hiroshi Sakai (Ehime University) for his support in the RNAscope analysis and Shin Fujimaki (Kumamoto University) for helping with single-cell and -nucleolus RNA-seq analysis of their data. This work was supported by grants from the MEXT Leading Initiative for Excellent Young Researchers (R.F.), Grant-in-Aid for Early-Career Scientists (21K17679; R.F.), and Grant-in-Aid for Scientific Research (B) (24K02876; R.F.) from the Japan Society for the Promotion of Science (JSPS). This research was also supported by AMED-CREST (JP23gm171008h). This work was also supported, in part, by grants to R.F. from the Takeda Science Foundation and Mochida Memorial Foundation for Medical and Pharmaceutical Research. Furthermore, this work received support from the Grant-in-Aid for the Japan Aerospace Exploration Agency (14YPTK-407 005512; S.T.) and the Grant-in-Aid for Scientific Research on Innovative Area from MEXT (18H04965; S.T.). T.H. and S.S were funded by JSPS Research Fellowships for Young Scientists (22J10954 [T.H.] and 23KJ0287 [S.S.]).

## Competing interests

The authors declare no competing financial interests.

## Author contributions

T.H., S.M., S.T., and R.F. conceived the study and designed the experiments. T.H., S.S., R.T., R.O., S.F., M.K., A.N., Y.O., M.M., S.M., D.S., E.W., T.K., and R.F. performed the experiments and bioinformatics analysis. T.H., G.W., T.S., and R.F. performed single-cell RNA-seq analysis. T.H., S.T., and R.F. wrote the manuscript. All authors contributed to the analysis and interpretation of results and reviewed the manuscript.

## Data Availability Statement

- All unique reagents generated in this study are available from the corresponding author, Ryo Fujita, in accordance with the relevant material transfer agreements.
- Single-cell RNA-seq data have been deposited in the NCBI Gene Expression Omnibus (GEO) and can be accessed with GSE250228.
- This paper does not report original code.
- Any additional information required to reanalyze the data reported in this paper is available from the corresponding author, Ryo Fujita upon request.

## Additional information

Supplementary Information is available for this paper.

## Notes

### Competing Interest Statement

The authors have declared no competing interest.

### Summary of Updates

We present new experimental evidence showing that the transition from proliferation to differentiation in muscle stem cells (MuSCs) is driven by Dusp13 and Dusp27. Additionally, we conducted further experiments that reinforce our conclusion that MYOD, rather than PAX7 or MYOGENIN, regulates Dusp13 and Dusp27.

