## Supplementary Information for "Dual-specificity phosphatases 13 and 27 as key switches in muscle stem cell transition from proliferation to differentiation"

### Supplementary Data

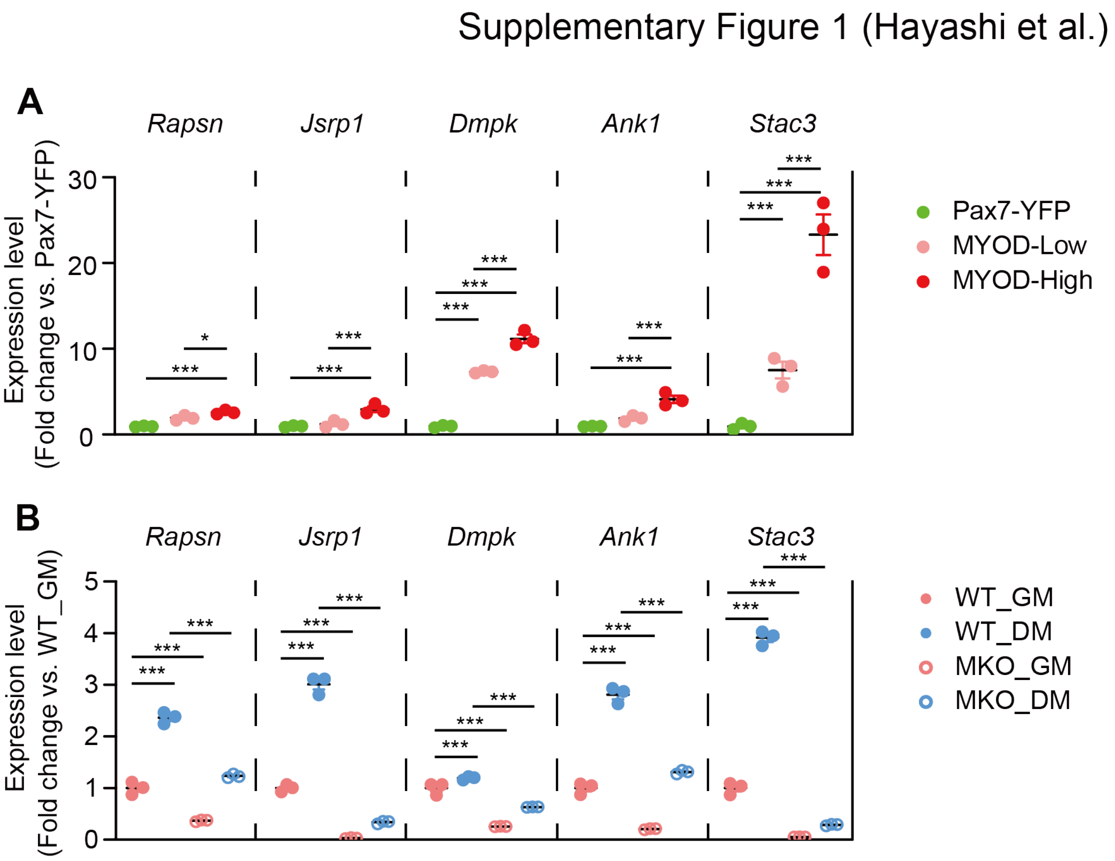


**Figure S1. Expression changes of candidate genes that are potentially MyoD-regulated.**

**(A)** Gene expression of *Rapsn*, *Jsrp1*, *Dmpk*, *Ank1*, and *Stac3* in RNA-seq data from Pax7-YFP (Quiescent MuSCs; QSCs), MyoD-low (MyoD-tdTomato low), and MyoD-high (MyoD-tdTomato high) (n = 3 per group). **(B)** Gene expression of *Rapsn*, *Jsrp1*, *Dmpk*, *Ank1*, and *Stac3* in RNA-seq between wild-type (WT)_growth medium (GM) (WT_GM), *MyoD* knock-out_GM (MKO_GM), WT_differentiation medium (WT_DM), and MKO_DM (n = 3 per group).

All data are represented as the mean ± standard error of the mean (SEM)

P values calculated using edge R **(A)** (FDR-corrected), DEseq2 **(B)** (FDR-corrected); *P < 0.05, ***P < 0.001 among all groups.

Supplementary Figure 2 (Hayashi et al.)

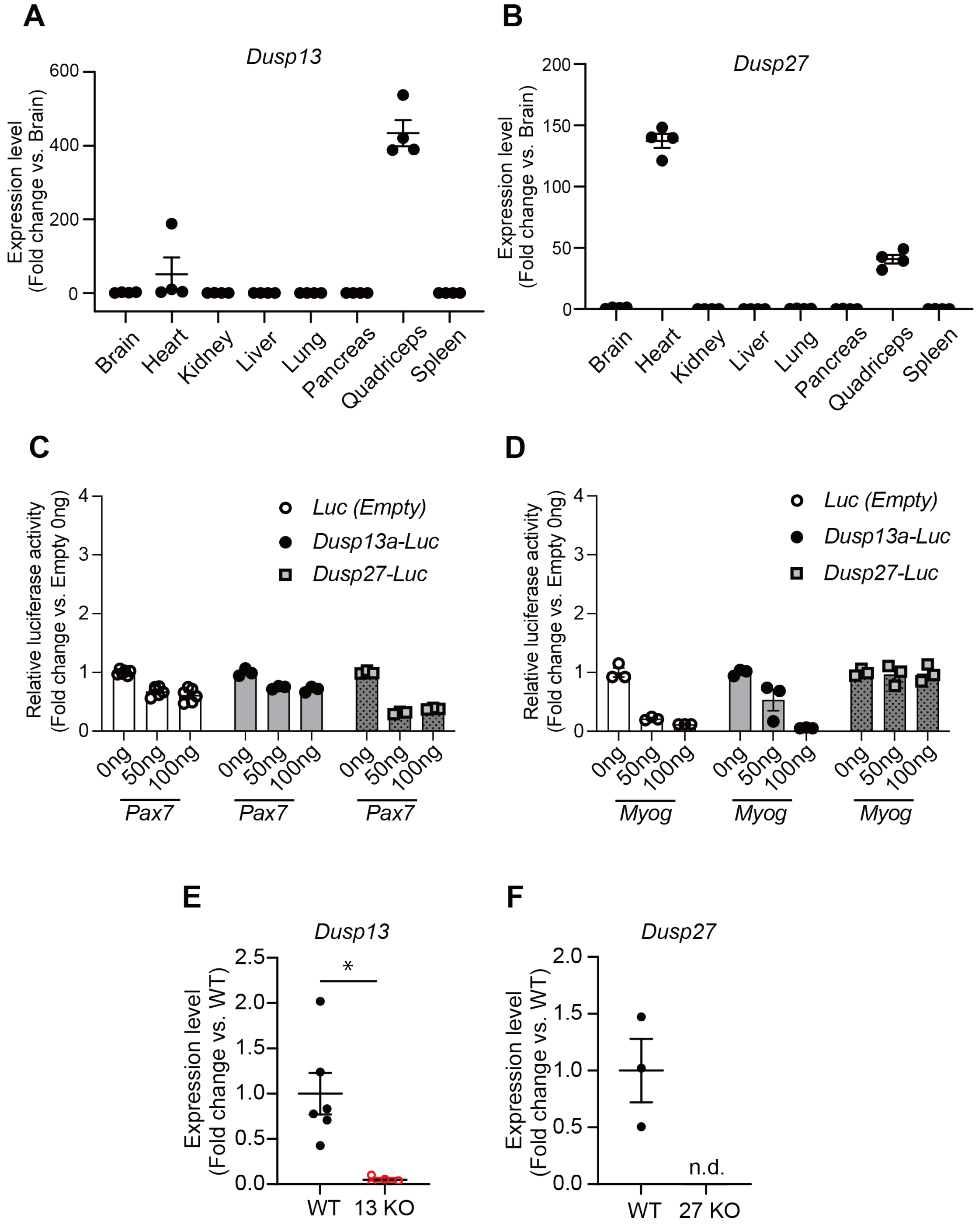

**Figure S2. Analysis of Dusp13 and Dusp27 expression across tissues and the influence of Pax7 and Myogenin on their promoter activity.**

**(A, B)** qPCR analysis of *Dusp13* (A) and *Dusp27* (B) mRNA levels in the tissues (brain, heart, kidney, liver, lung, pancreas, muscle, and spleen) of WT mice at 13–14 weeks (n = 4 per tissue).

**(C, D)** Relative *Dusp13*-luciferase reporter or *Dusp27*-luciferase reporter activities in HEK293T cells overexpressing *Pax7* **(C)** and *Myogenin* **(D)** (n = 3 per group).

**(E)** qPCR analysis of *Dusp13* mRNA levels in the gastrocnemius muscles of WT and *Dusp13* KO (13 KO) mice at 8–12 weeks (n = 4–6 independent experiments).

**(F)** qPCR analysis of *Dusp27* mRNA levels in the gastrocnemius muscles of WT and *Dusp27* KO (27 KO) mice at 8–12 weeks (n = 3 per group).

All data are represented as the mean ± standard error of the mean (SEM)

P values calculated using Student's t-test; *P < 0.05, n.d., not detected.

Supplementary Figure 3 (Hayashi et al.)

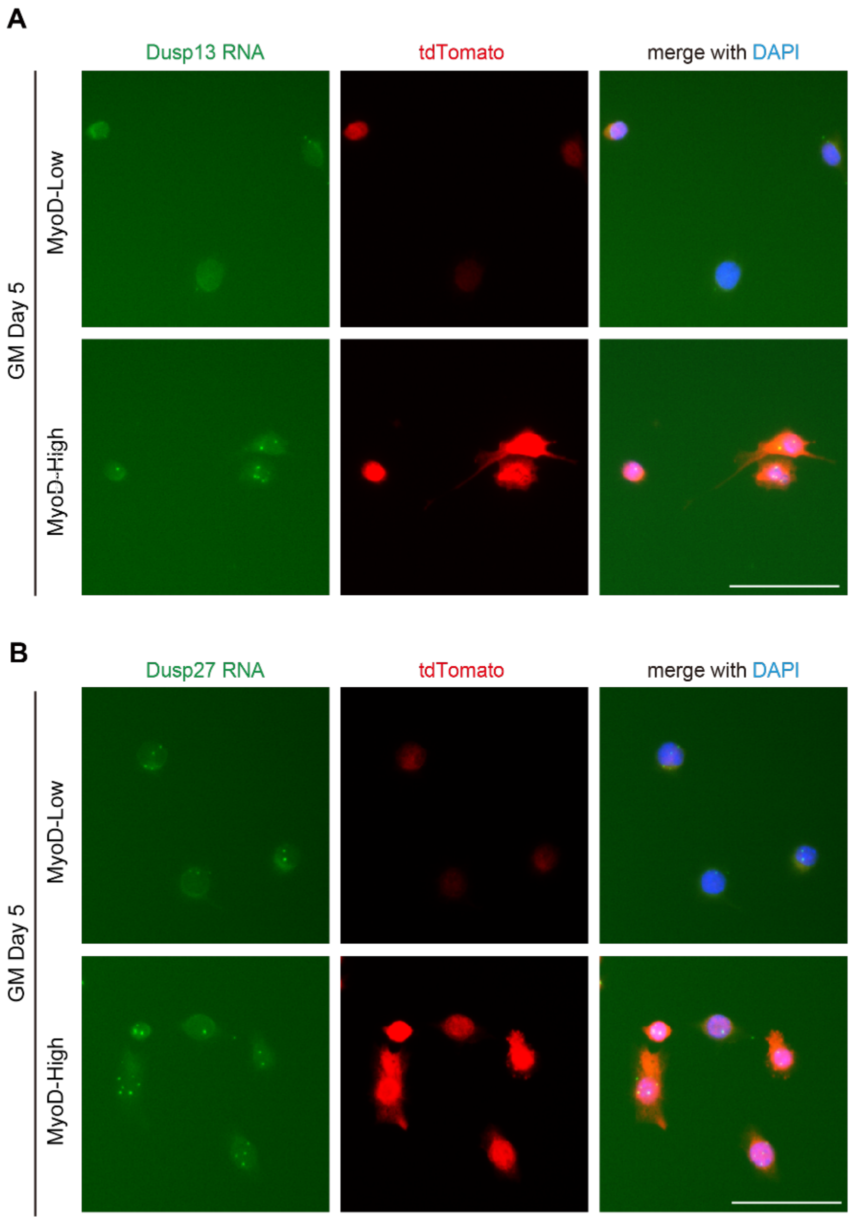


**Figure S3. Detection of Dusp13 and Dusp27 mRNAs in MyoD-low and MyoD-high MuSC by RNAscope**
**(A, B)** RNAscope of MyoD-low and MyoD-high MuSCs cultured for 5 days in GM probed for *Dusp13* **(A, green)** and *Dusp27* **(B, green)**. After hybridization, MyoD-tdTomato was visualized using an antibody against tdToamto (red). Nuclei are stained with DAPI. Scale bar: 100 µm. Scale bar: 50µm.

Supplementary Figure 4 (Hayashi et al.)

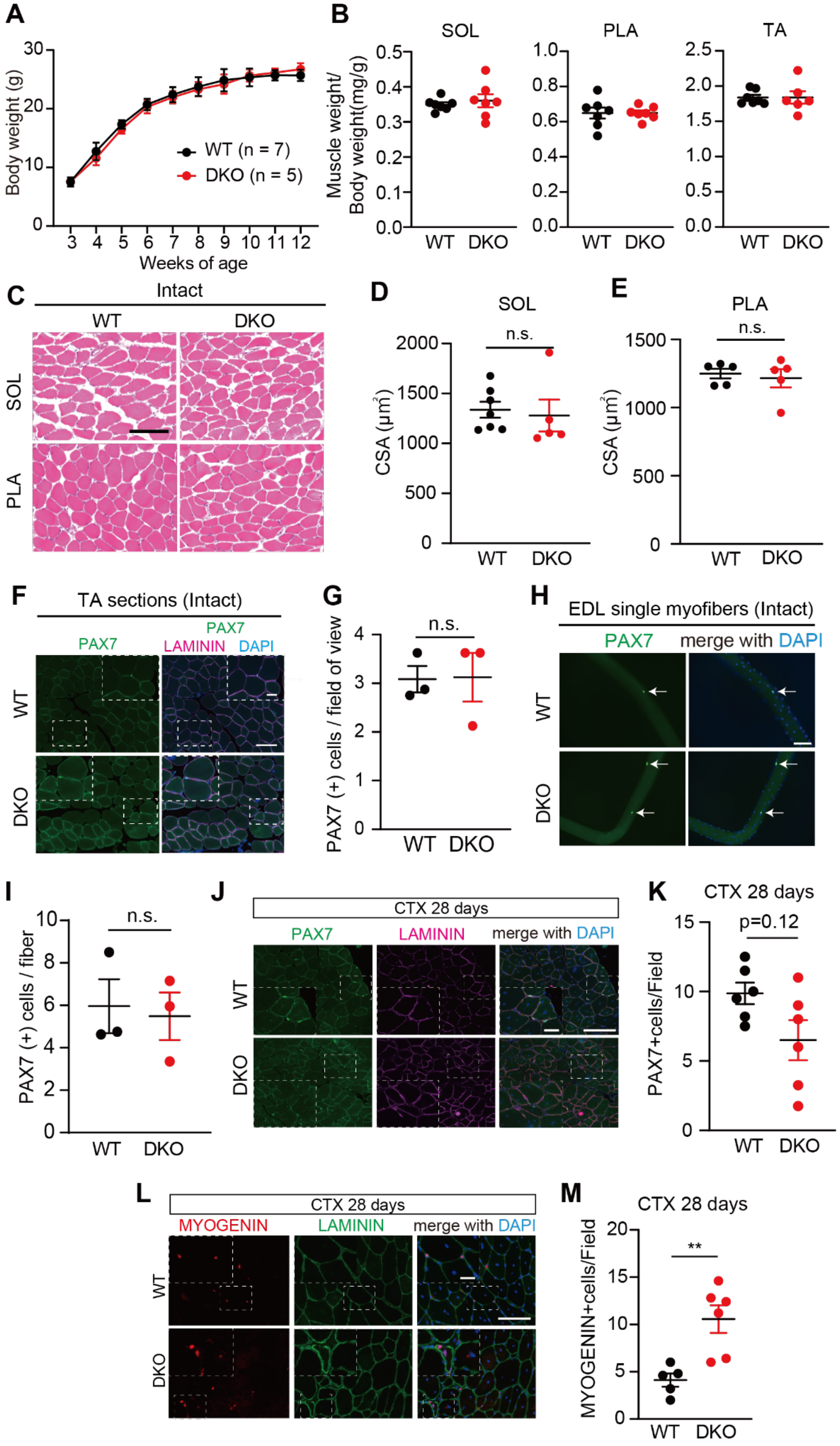

**Figure S4. DKO mice exhibit normal muscle development but show delayed muscle differentiation at day 28 post-injury.**

**(A)** Body weight changes of male *Dusp13*:*Dusp27* double knock-out (DKO) and WT mice at 3 to 12 weeks of age (n = 5–7 mice per group).

**(B)** Skeletal muscle weights of male DKO (n = 6) and WT (n = 7) mice at 13 weeks of age, normalized to body weight.

**(C)** H&E staining of SOL and PLA muscle cross-sections. Scale bar: 100 μm.

**(D, E)** CSA of SOL **(D)** or PLA **(E)** muscles of male DKO and WT mice at 13 weeks (n = 5–7 independent experiments).

**(F)** Immunoflurescence of PAX7 and LAMININ in intact TA muscles. Scale bar: 100 μm. The area in the white dotted boxes is shown at a higher magnification. Scale bar: 25 μm.

**(G)** Average number of PAX7 (+) nuclei in DAPI (+) in **(F)**.

**(H)** Immunofluorescence of PAX7 on freshly isolated single extensor digitorum longus (EDL) myofibers from DKO and WT mice at 13 weeks (n = 3 per group). Nuclei stained with DAPI. White arrows indicate PAX7 (+) cells on EDL myofibers. Scale bar: 100 μm.

**(I)** Number of PAX7 (+) cells EDL myofibers in **(H)**.

**(J)** Immunofluorescence of PAX7 with LAMININ on sections of TA muscles from WT and DKO mice, 28 days post-CTX injury. Nuclei are stained with DAPI. Scale bar: 100 μm. The area in the white dotted boxes is shown at a higher magnification. Scale bar: 25 μm.

**(K)** Average number of PAX7 (+) nuclei in **(J)** are quantified (n = 6 per group).

**(L)** Immunofluorescence of MYOGENIN with LAMININ on sections of TA muscles from WT and DKO mice, 28 days post-CTX injury. Nuclei are stained with DAPI. Scale bar: 100 μm. The area in the white dotted boxes is shown at a higher magnification. Scale bar: 25 μm.

**(M)** Average number of MYOGENIN (+) nuclei in **(L)** are quantified (n = 4–6 independent experiments).

All data are represented as the mean ± standard error of the mean (SEM)

P values calculated using Student's t-test; $^{**}P < 0.01$, n.s., not significant.

### Supplementary Figure 5 (Hayashi et al.)

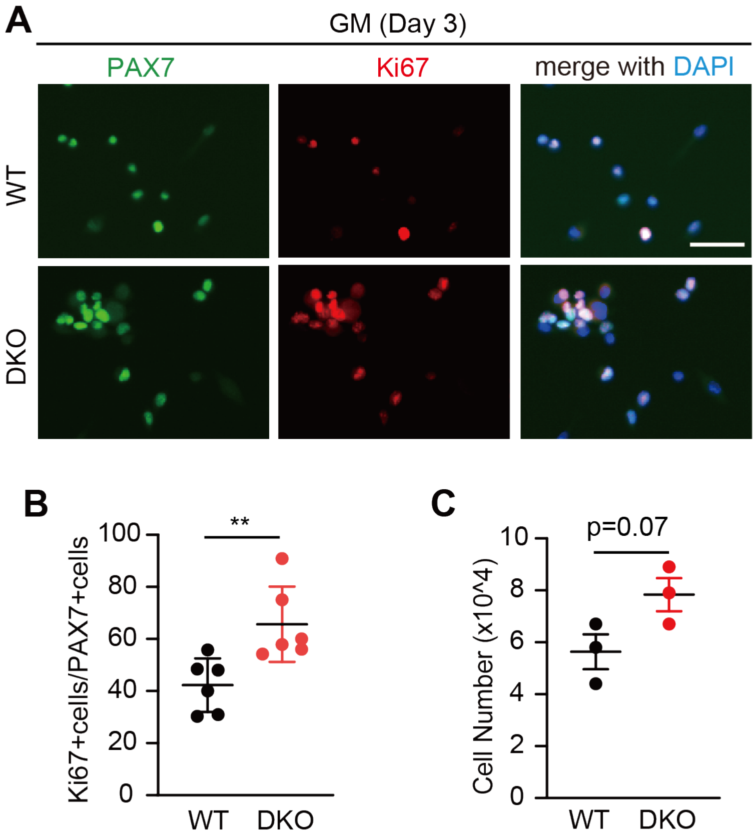


**Figure S5. MuSCs derived from *Dusp13*:*Dusp27* DKO mice stay in the proliferation stage.**

**(A)** Immunofluorescence of PAX7 and Ki67 on MuSCs cultured for three days in a growth medium (GM).

**(B)** Average number of Ki67 (+) nuclei in PAX7 (+) in **(A)** are quantified (n = 6 per group).

**(C)** Cell numbers of MuSCs derived from WT or DKO at GM day 4 (n = 3 per group).

All data are represented as the mean ± standard error of the mean (SEM)

P values calculated using Student's t-test; **P < 0.01.

Supplementary Figure 6 (Hayashi et al.)

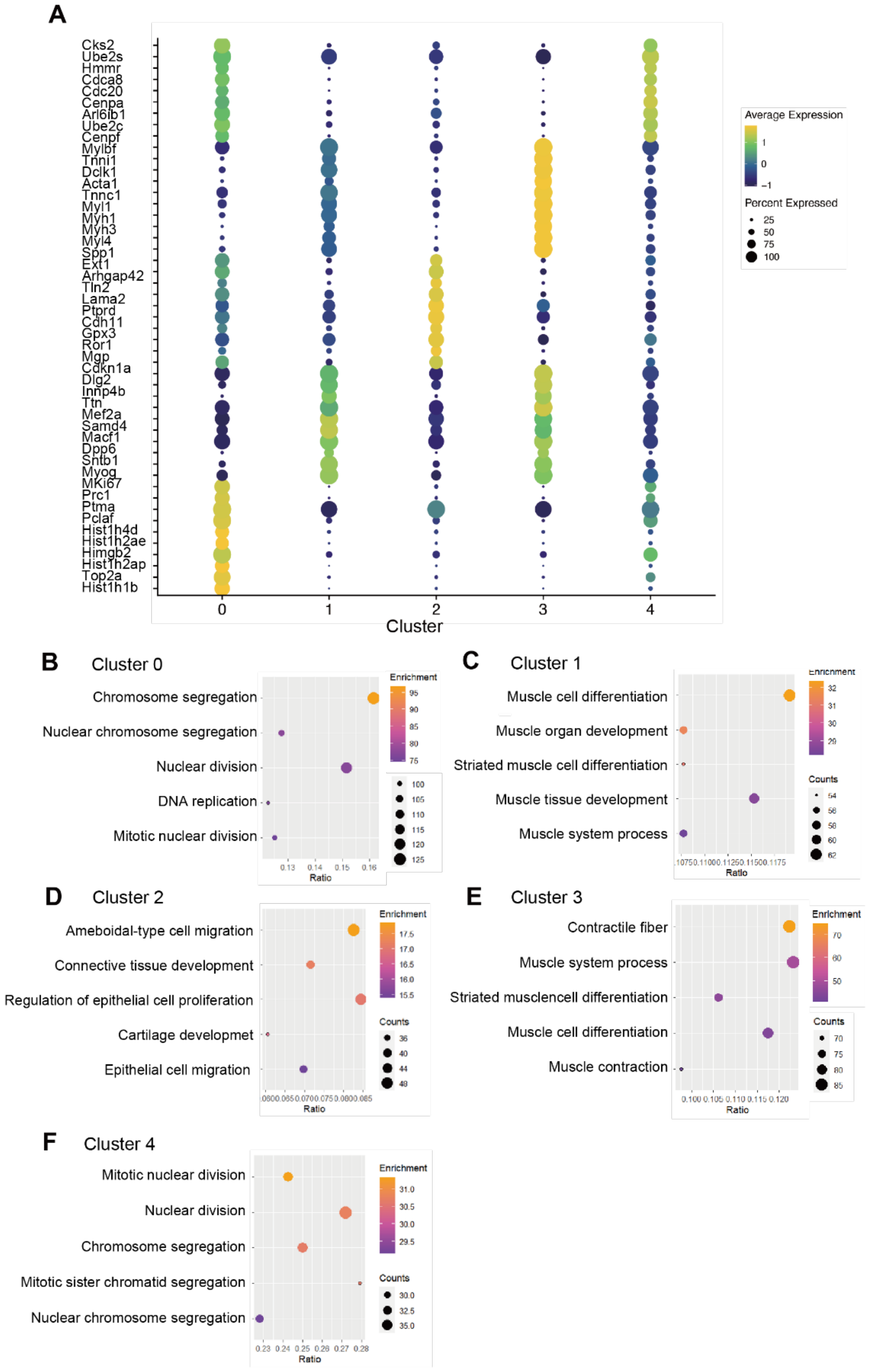


**Figure S6. Pathway analysis in single-cell RNA-sequencing data of MYOD-high MuSCs from MyoD-KI mice.**

**(A)** Dot plot representing the expression of the top 10 differentially expressed genes for each cluster.

**(B–F)** A Gene Ontology analysis for cluster 0 **(B)**, cluster 1 **(C)**, cluster 2 **(D)**, cluster 3 **(E)**, and cluster 4 **(F)**.

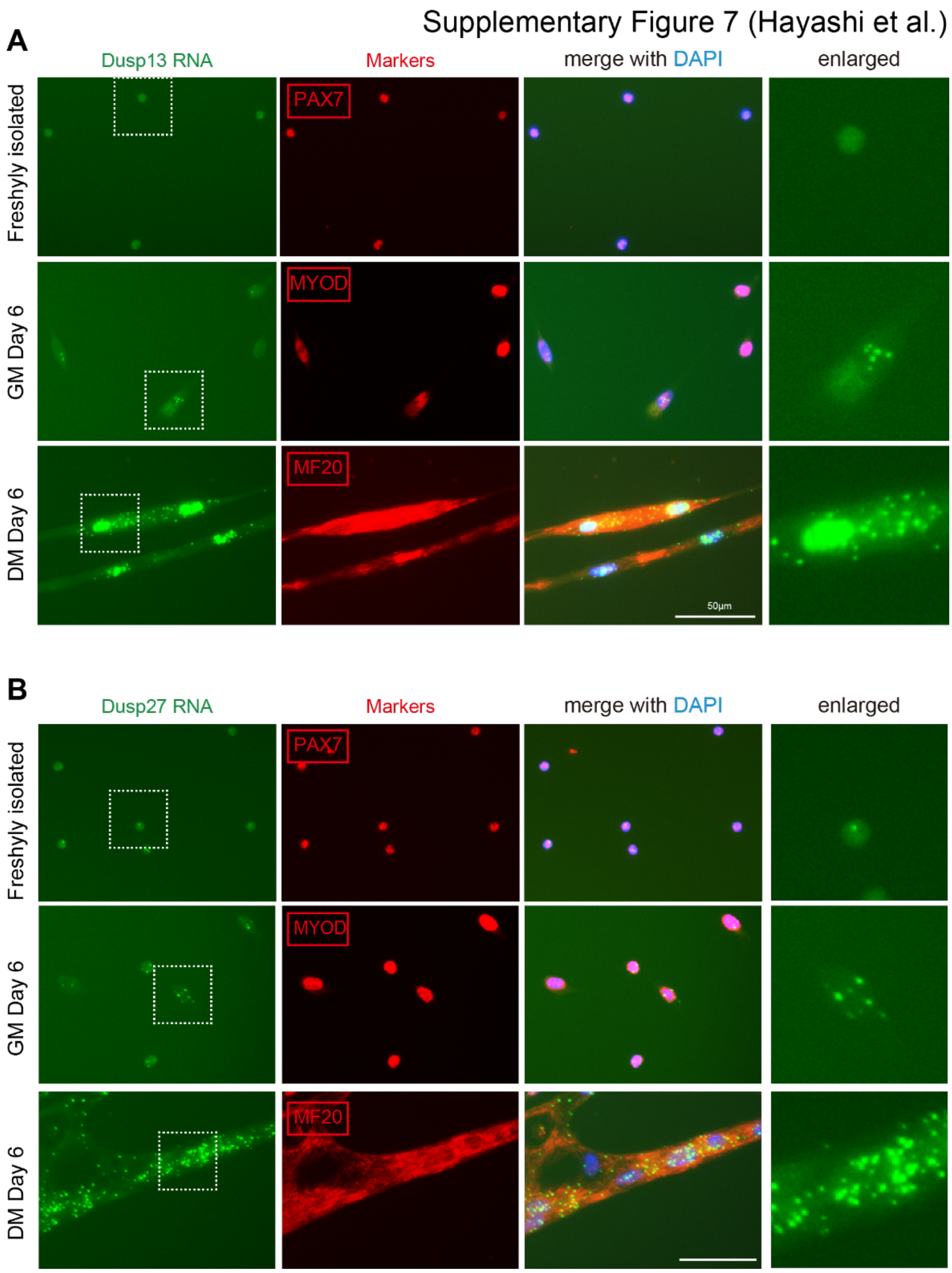


**Figure S7. Detection of Dusp13 and Dusp27 mRNAs in freshly isolated, cultured MuSCs, and myotubes. (A, B)** RNA-scope to validate the expression of *Dusp13* mRNA **(A, green)** and *Dusp27* mRNA **(B, green)** in freshly isolated MuSCs (cultured for 1h for attachment to the dish), activated MuSCs for 6 days in growth medium (GM), and myotubes cultured for 6 days in differentiation medium (DM). In both conditions, immunostaining for marker proteins at different stages of MuSCs was performed using antibodies against, PAX7, MYOD, and MF20 after hybridization. Nuclei are stained with DAPI. Scale bar = 50 µm.

Supplementary Figure 8 (Hayashi et al.)

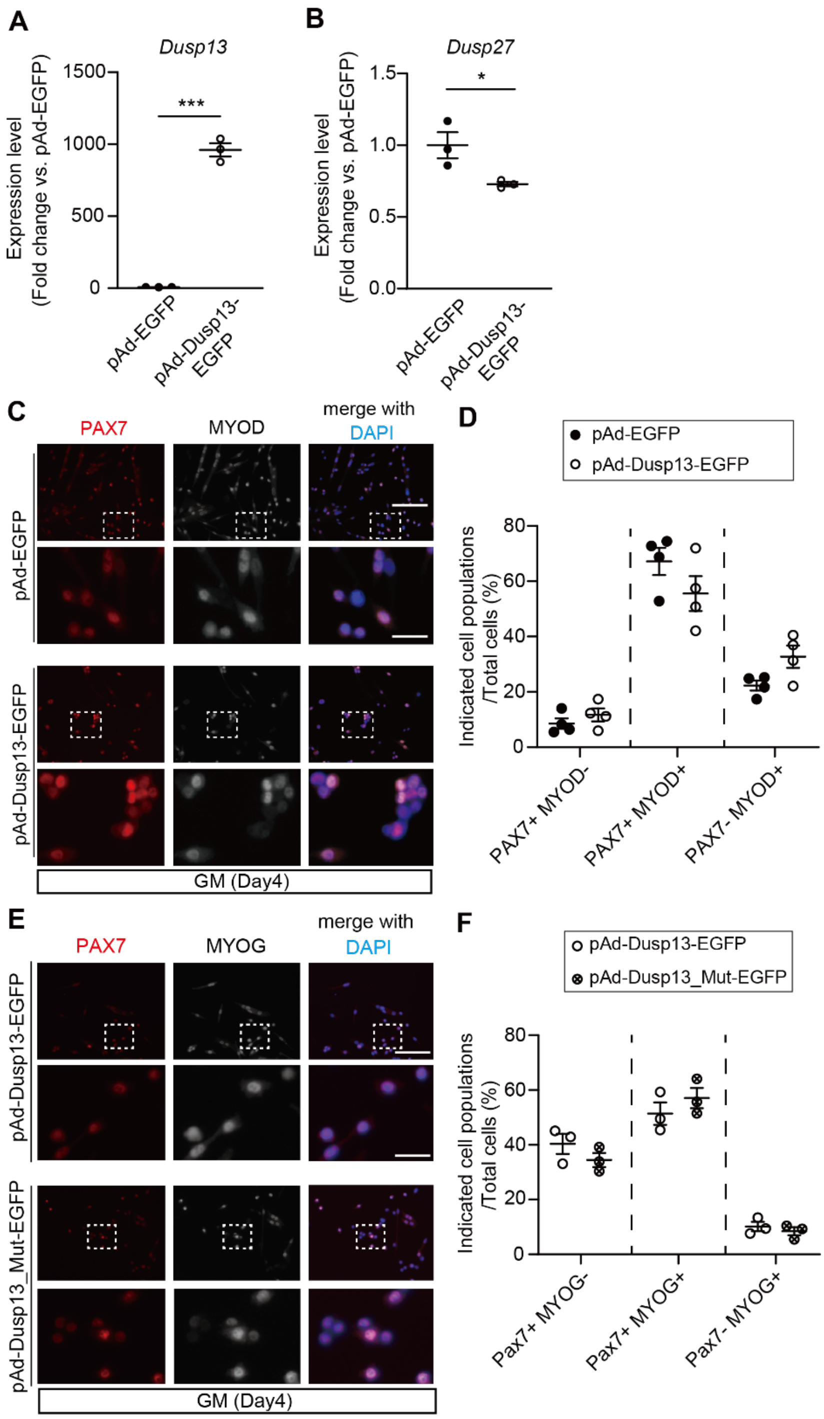


**Figure S8. Expression levels of *Dusp13* and *Dusp27* in a *Dusp13* overexpression experiment.**

**(A, B)** qPCR analysis of *Dusp13* **(A)** and *Dusp27* **(B)** mRNA levels in MuSCs which was overexpressed EGFP or *Dusp13*-EGFP (n = 3 per group).

**(C)** Immunofluorescence of PAX7 and MYOD on cultured MuSCs after transduction with adenovirus vectors overexpressing EGFP or Dusp13-EGFP. Nuclei are stained with DAPI (n = 3 per group).

**(D)** The proportion of each population in (**C**).

**(E)** Immunofluorescence of PAX7 and MYOGENIN on cultured MuSCs after transduction with adenovirus vectors overexpressing Dusp13 (pAd-Dusp13-EGFP) or Dusp13 without phosphatase activity (pAd-DUSP13_Mut-EGFP). Nuclei are stained with DAPI (n = 3 per group).

**(F)** The proportion of each population in (**E**).

All data are represented as the mean ± standard error of the mean (SEM)

P values calculated using Student's t-test; *P < 0.05, ***P < 0.001.

**Table S1. List of oligonucleotide sequences used as primers**

| Gene (genotyping) | Orientation | Sequence |
|---|---|---|
| *Dusp13* (WT) | Fw | 5′-GAATGAGGCTATTACCCGGC-3′ |
| | Rv | 5′-ACATAACCCACTCCTTCCGG-3′ |
| *Dusp13* (KO) | Fw | 5′-CCCACATACATATTTGTAGGCACT-3′ |
| | Rv | 5′-GCCCCAATGCTAGGTTTACA-3′ |
| *Dusp27* (WT) | Fw | 5′-AACCGAAAGCATCTTCATGG-3′ |
| | Rv | 5′-GTGGGCTAGAGAAGCATTCG-3′ |
| *Dusp27* (KO) | Fw | 5′-TTGACCTGCCCTACATCTCC-3′ |
| | Rv | 5′-CTCACCCCAGGACAAGATGT-3′ |
| Gene (RT-qPCR) | Orientation | Sequence |
| *b-actin* | Fw | 5′-CATTGCTGACAGGATGCAGAAGG-3′ |
| | Rv | 5′-TGCTGGAAGGTGGACAGTGAGG-3′ |
| *Dusp13* | Fw | 5′-AAGTCTGGCCCAACCTTTTC-3′ |
| | Rv | 5′-CACTGCTGCCGTAGAAGTCA-3′ |
| *Dusp27* | Fw | 5′-AACCGAAAGCATCTTCATGG-3′ |
| | Rv | 5′-AGCCACGCTTTTCTCAGCTA-3′ |
| Gene (Cloning) | Orientation | Sequence |
| *Dusp13* for Adenovirus | Fw | 5′-CCCAGGATCCATGTCACAGACC-3′ |
| | Rv | 5′-GGGTCTCGAGCTAGTGCTGCTGGCTC-3′ |
| *Dusp13* for Luciferase | Fw | 5′-GCTCGCTAGCCTCGATAACCCTTGCAACAACTCC-3′ |
| | Rv | 5′-TCTTGATATCCTCGAACTTCATCCACTCTACTACA-3′ |
| *Dusp27* for Luciferase | Fw | 5′-GCTCGCTAGCCTCGAATGTCTCCCACGAAGCCT-3′ |
| | Rv | 5′-TCTTGATATCCTCGAACACTCAGGGCAGACTGG-3′ |
| *MyoD* | Fw | 5′-GGATTCGCGAGAATTATGGAGCTTCTACGCCG-3′ |
| | Rv | 5′-CGACTCTAGAGAATTTCAAAGCACCTGATAAATCG-3′ |
| *Pax7* | Fw | 5′-CTCCCCAGGGGGATCATGGCGGCGCTGCCCGGC-3′ |
| | Rv | 5′-GAGGTTGATTGTCGACTAGTAGGCTTGTCCCGTTTCC-3′ |
| *Myogenin* | Fw | 5′-CIGGCGGCCGCTCGAGATGGAGCTGTATGAGACATC-3′ |
| | Rv | 5′-TGCTCACCATCTCGAGGTTGGGCATGGTTTCGTC-3′ |
| Gene (Mutation) | Orientation | Sequence |
| *Dusp13 D97A* | Fw | 5′-GACCCCTAGGGTCGGGTGCGGGAGGGA-3′ |
| | Rv | 5′-CTATAACTTTAGTCCCTCCCGCACCCG-3′ |
| *Dusp13 C128S* | Fw | 5′-CGATTCCACGACCACGTGAGGCACCAC-3′ |
| | Rv | 5′-AGCCGAGTGTGGGTGGTGCCTCACGTG-3′ |
